# Symbiotic entrenchment through ecological Catch-22

**DOI:** 10.1101/2025.07.02.662832

**Authors:** Thomas H. Naragon, Joani. W. Viliunas, Mina Yousefelahiyeh, Adrian Brückner, Julian M. Wagner, Hannah M. Ryon, Danny Collinson, Sheila A. Kitchen, Reto S. Wijker, Alex L. Sessions, Joseph Parker

## Abstract

Symbiotic organisms frequently evolve obligate dependencies on hosts, but the evolutionary changes that entrench such lifestyles are poorly understood. Ant societies are vulnerable to parasitic “myrmecophiles”: impostor species that infiltrate colonies and are often unable to survive outside of them. Here we show that obligate dependence of a myrmecophile on its host arises from irreversibility of the fundamental steps that achieve social acceptance inside the nest. We report a convergent system in which parallel rove beetle lineages (Staphylinidae) evolved from free-living ancestors to parasitize the same host ant. Exploiting this system, we uncover cellular mechanisms by which these beetles mimic host ant cuticular hydrocarbons (CHCs): nestmate recognition pheromones, which function pleiotropically to prevent desiccation. We present evidence of a biological stealth mechanism in a rove beetle in which the CHC biosynthetic machinery becomes transcriptionally silenced on entering the nest. Silencing transforms the beetle into a chemical blank slate onto which ant CHCs are horizontally transferred via interspecies grooming behavior. This strategy leads to identical chemical resemblance and seamless social integration within the colony. CHC pathway silencing is irreversible, however, forcing the beetle into a chronic, physically close dependence on ants to both maintain nestmate status and prevent desiccation. Loss of CHC silencing renders the beetle detectable to ants; conversely, loss of behavioral attraction to ants renders the beetle desiccation prone. Our findings show how symbiotic entrenchment can arise from a Catch-22-like ratchet operating at the organismal level.

---

Obligate symbioses are commonly viewed as embodying irreversible evolution. Diverse animal^1,2^, plant^3,4^, fungal^5,6^ and microbial^7,8^ clades spanning the spectrum of mutualism to parasitism exist in which reversion to the free-living state is unknown, or extremely rare^9–14^. The loss of traits^15,16^ and genes^17–38^ that facilitate free-living lifestyles is pervasive in symbionts—the outcome of relaxed selection in host environments^28,37–44^, and gene-inactivating mutations that prove adaptive in specialized niches^36,45–50^. The accumulation of reductive changes is a process widely thought to counteract evolutionary reversion from different forms of symbiosis^23,25,51–53^. Yet, whether the loss of traits and genes is the primary cause of entrenchment, as opposed to a reinforcing consequence, is challenging to disentangle. The antiquity of many symbiotic lineages obscures the coercive forces that drove free-living ancestors towards obligate symbiosis. During this transition, critical changes in cellular, physiological or behavioral mechanisms arose that locked the phenotype into host dependence. The often-highly modified genomes and derived phenotypes of extant symbionts mask what these steps were; how they conspired to obstruct reversion to the free-living state is consequently unknown for symbiotic taxa across phylogeny.

Myrmecophiles are symbiotic organisms specialized for life inside ant societies^54–56^. Approximately 100,000 animal species are suspected to engage in this form of symbiosis^57^, including many that obligately depend on host colonies and are unable to live away from them. Rove beetles of the subfamily Aleocharinae comprise a megadiverse clade in which dozens of lineages have convergently evolved from free-living predators into specialized myrmecophiles that obligately assimilate into host colony social structure^55,58–61^. The transition to colony life involves a radical transformation of the phenotype, encompassing stereotyped innovations in behavior, chemical ecology and anatomy that enable aleocharine myrmecophiles to manipulate host ants and gain social acceptance within the nest. These intimate relationships contrast with free-living species that display defensive behavior towards ants—the ancestral condition in Aleocharinae^55,62–65^. Myrmecophile clades with the most extreme symbiotic phenotypes appear seamlessly integrated into colonies; within these groups, there are no apparent cases of reversion to the free-living state^60,61^. This repeated trend implies entrenchment may be inherent to the assimilation of aleocharine lineages into host ant societies. Here, we leverage this convergent system to retrace how evolving into a host niche can be intrinsically irreversible, locking a lineage into perpetual symbiosis.

## A convergent system of ant-myrmecophile symbiosis

We identified a natural system of ant-myrmecophile symbiosis in Southern California in which three lineages of aleocharine rove beetle have converged to socially parasitize the same host ant, the velvety tree ant (*Liometopum occidentale*)^66^. Time-calibrated phylogenomic analysis reveals that the three beetles—*Platyusa sonomae*, *Sceptobius lativentris* and *Liometoxenus newtonarum*—belong to distinct aleocharine tribes (Lomechusini, Sceptobiini and Oxypodini, respectively) and share a common ancestor ∼89 mega-annum (Ma, 95% HPD: 106-74 Ma) ago that was likely free-living (**Fig. 1a; Fig. S1a, b**). Each lineage separately transitioned to myrmecophily, with inferred divergence times from free-living sister groups of 25.5 Ma (*Platyusa,* 95% HPD: 36-15 Ma), 30.8 Ma (*Sceptobius,* 95% HPD: 46-17 Ma), and 51 Ma (*Liometoxenus,* 95% HPD: 70-30 Ma) (**Fig. 1a; Fig. S1a, b**). These times approximate or postdate the inferred age of the genus *Liometopum* within its ant subfamily, Dolichoderinae^67^, but represent maximum age estimates. Each transition likely occurred still more recently, on account of probable sister taxa known to us that are phylogenetically even closer than those included in our cladogram.

**Figure 1:**
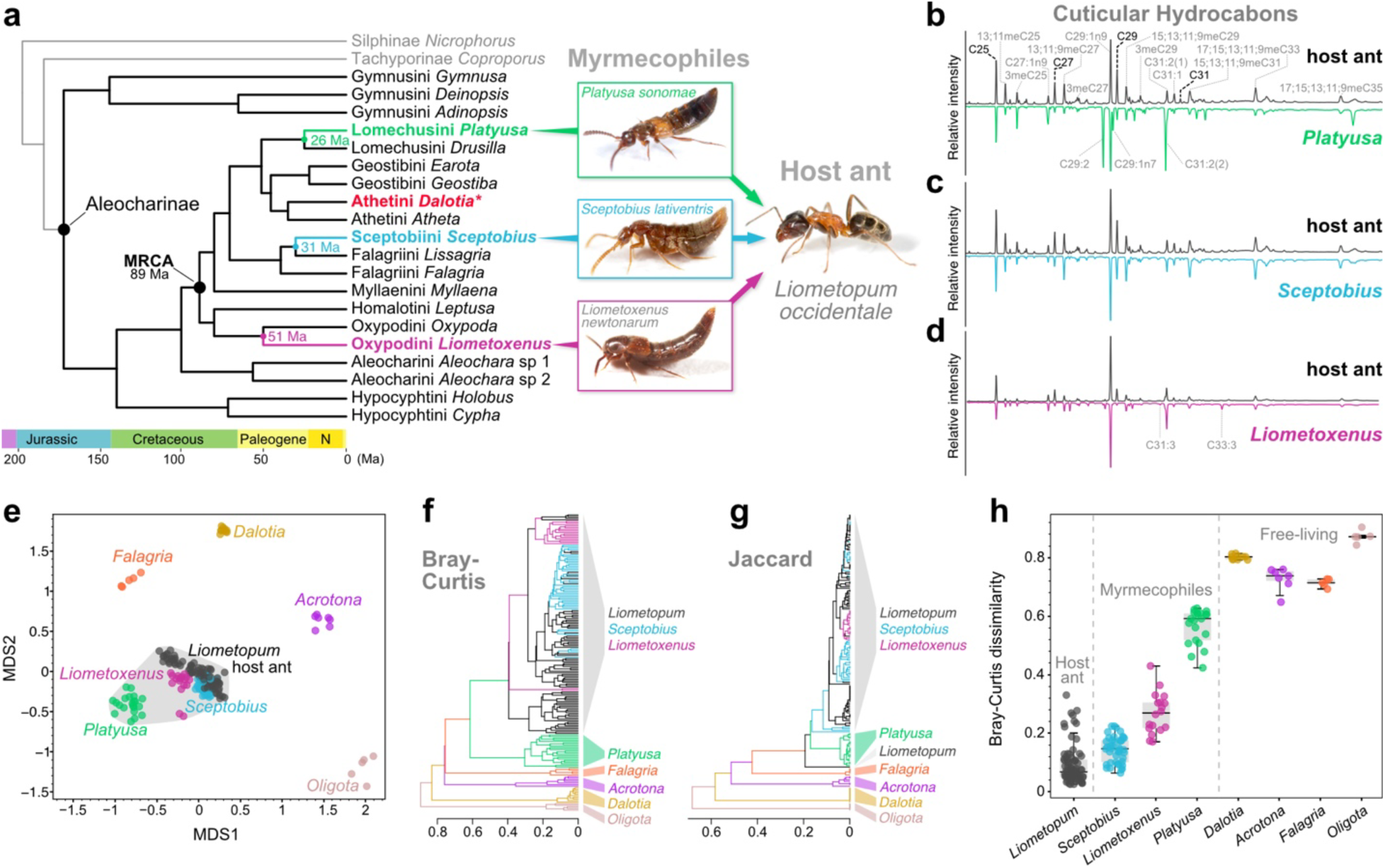
A convergent system of ant-myrmecophile symbiosis. **a:** Dated phylogenomic tree inferred from 1039 orthologous protein-coding loci and 12 fossil calibration points. The topology is strongly supported at all nodes (**Fig. S1b)**. **b-d:** Gas chromatographs of CHCs from *Liometopum* host workers (top trace, with identified compounds labelled), and the myrmecophiles *Platyusa* (**b**), *Sceptobius* (**c**) and *Liometoxenus* (**d**). Unique compounds absent from the ant trace are labelled for Platyusa and Liometoxenus. **e:** NMDS ordination of CHC chemical dissimilarity (pairwise Bray-Curtis) between the host ant and three myrmecophiles (demarcated by grey convex hull), and four free-living aleocharine species, 2D stress = 0.13. **f, g:** Hierarchical clustering of individual insect CHC profiles based on compound identity and relative abundance (Bray Curtis) (**f**), and identity alone (Jaccard) (**g**). **h:** Bray Curtis dissimilarity of individual ants and myrmecophiles to the mean ant profile of their colony of origin; dissimilarities of free-living beetles are to the mean ant profile calculated across colonies.

## Convergent evolution but varying accuracy of CHC mimicry

Colonies of *Liometopum occidentale* are vast, comprising ∼10^6^ workers that are highly aggressive towards other organisms^66^. Our field data obtained over eight years, plus historical collection records^68–70^, indicate all three beetles have been found exclusively with *Liometopum* ants, either inside colonies, at nest entrances, or along foraging trails. To understand how these myrmecophiles evade detection by their host, we examined one of the primary communication systems used by ants. Cuticular hydrocarbons (CHCs) are very long-chain alkanes and alkenes that many insects use as contact pheromones^71–73^. CHCs are synthesized in abdominal cells named oenocytes^74^ and are secreted onto the body surface, forming a waxy pheromonal coating that is essential for desiccation avoidance^75–78^. Ants have co-opted CHCs as nestmate recognition cues, producing a complex CHC cocktail that functions as a colony identifier. Genetic and environmental factors cause the CHC template to vary between colonies, and temporally within the same colony^79^. Via antennal detection, workers evaluate CHCs on the bodies of other insects, accepting nestmates that share the prevailing colony template, while aggressing insects that present divergent profiles^72,79,80^.

Gas chromatography-mass spectrometry (GCMS) analysis of body surface compounds revealed that the three myrmecophiles present CHC profiles strikingly similar to that of *Liometopum* workers (**Fig. 1a-c)**. We identified each host ant CHC and found that all three myrmecophiles likewise possess the same compounds without exception (**Table S1**). Notably, *Liometoxenus* and *Platyusa* — but not *Sceptobius —* possess a small number of additional CHCs (all alkenes) that are absent from the ant’s profile (**Fig. 1b, d**; **Table S1**). We measured CHCs from many individuals of *Liometopum*, the three myrmecophiles, and four free-living species from across the Aleocharinae phylogeny: *Acrotona* and *Dalotia* (both tribe Athetini); *Oligota* (Hypocyphtini) and *Falagria* (Falagriini). Non-metric multidimensional scaling (NMDS) of pairwise Bray-Curtis dissimilarity measurements of the various profiles revealed that the three myrmecophile beetles cluster tightly with their host ant to the exclusion of the free-living beetles (**Fig. 1e**). Hierarchical clustering of pairwise Bray-Curtis or Jaccard dissimilarity measures further confirmed that the CHC profiles of the myrmecophiles and *Liometopum* were more similar to each other than to CHC profiles of free-living species, based on quantitative CHC composition (**Fig. 1f)** or qualitative compound presence/absence (**Fig. 1g**). We infer that the three myrmecophiles each independently evolved to chemically mimic their shared host.

CHC mimicry has been reported from diverse myrmecophiles^61,81–84^, and was likely a key step in the symbiotic evolution of *Platyusa*, *Sceptobius* and *Liometoxenus*, permitting closer contact with ants. The three beetles nevertheless differ in how accurately they match the ant’s profile. We quantified chemical divergence between beetle and ant on a nest-by nest basis, calculating the Bray-Curtis dissimilarity of every beetle to the mean profile of worker ants from that beetle’s host colony. Strikingly, *Sceptobius* beetles and host ants exhibit equivalent dissimilarity to the mean ant profile, indicating that ant and beetle have effectively indistinguishable CHC profiles (**Fig. 1h**). In contrast, the additional compounds in the profiles of *Liometoxenus* and *Platyusa* result in these species having significant dissimilarity to their host colony’s mean CHC profile; in the case of *Platyusa*, differences in the ratios of some compounds common to *Platyusa* and the host ant further increase dissimilarity (**Fig. 1h)**. Due to the sensitivity of ants to even minor deviations from the CHC template^85^, these dissimilarities are likely detectable by the ants, with consequences for the corresponding degree of integration of these myrmecophiles within the *Liometopum* society. Indeed, of the three beetles, *Sceptobius* frequents the nest proper^68^, where observations indicate it feeds on host eggs and larvae (**Video S1**). *Platyusa* and *Liometoxenus* by contrast are predominantly found at nest entrances and trails, and feed directly on adult workers (**Video S1**).

## Approximate chemical mimicry through *de novo* CHC synthesis

We investigated the mechanisms by which these myrmecophiles create their CHC profiles, focusing on *Platyusa* and *Sceptobius* that respectively exhibit the least and most accurate mimicry. We first determined the biosynthetic sources of the beetles’ CHCs, leveraging the tendency of carbon stable isotope ratios of specific molecules to vary between organisms depending on diet^86^. Using compound-specific gas chromatography isotope-ratio mass spectrometry (GC-IRMS), we measured the δ^13^C values of specific CHCs present on the two beetles and the ant. If a CHC is present on ants and beetles from the same colony but differs in δ^13^C per mille value between the two species, it independently originated in each species; conversely, if the CHC has an identical δ^13^C in both species, it probably came from the same biosynthetic source. We extracted CHCs from four replicates of twenty *Platyusa* beetles and twenty *Liometopum* ants from the same colony. GC-IRMS was able to achieve sufficient separation to measure δ^13^C of three compounds shared by both species: C25, C27 and C29. For all three CHCs, we determined a significant δ^13^C offset of ∼5 per mille between the *Platyusa* CHC and the corresponding compound from *Liometopum* (**Fig. 2A**). We deduce *Platyusa* synthesizes these hydrocarbons—and likely other CHCs in its profile—*de novo*.

**Figure 2.**
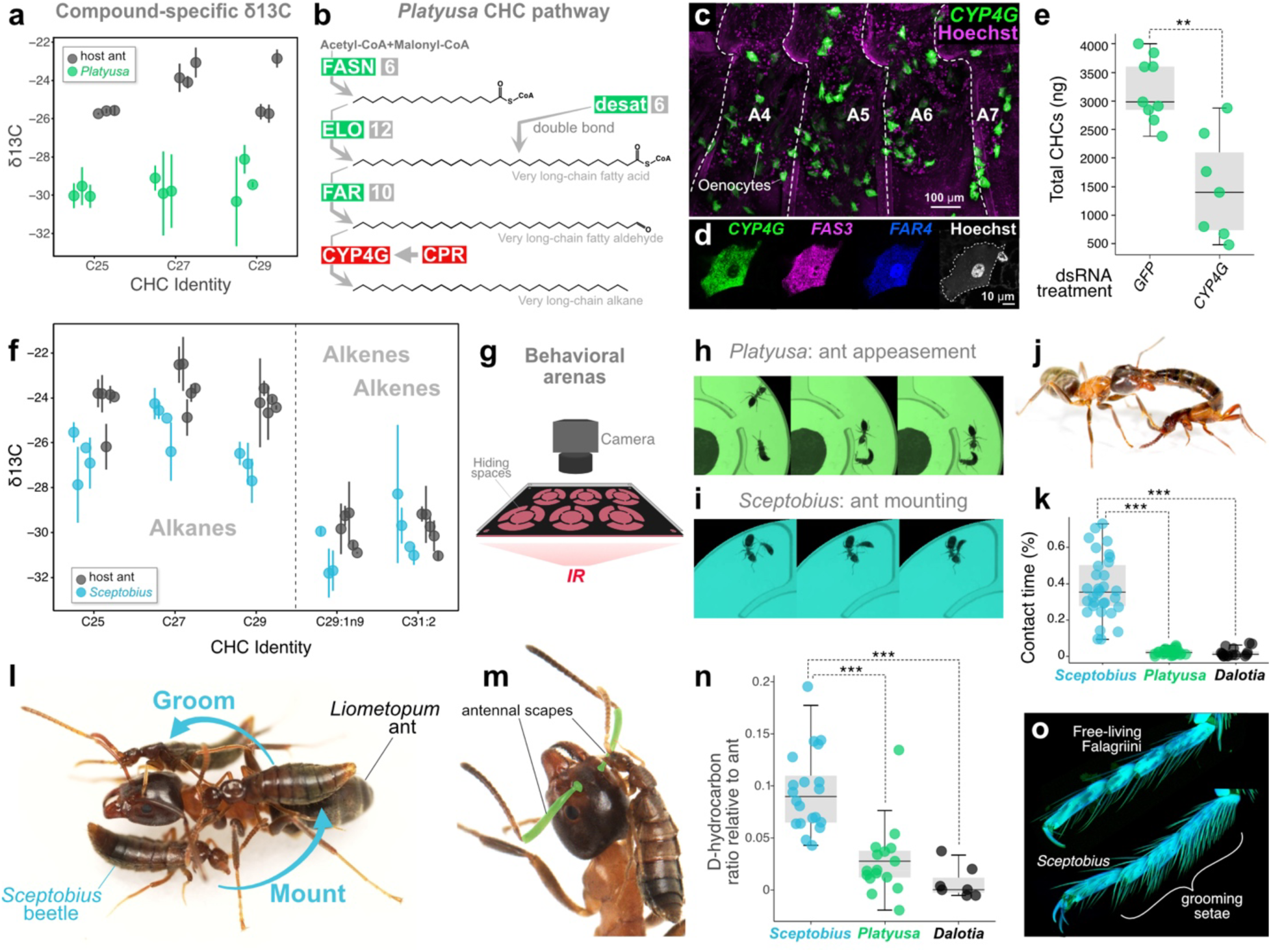
Distinct mechanisms of CHC mimicry. **a:** δC13 values of specific CHCs from *Platyusa* and *Liometopum* measured via GC-IRMS. **b:** The *Platyusa* CHC pathway. Grey boxes show number of paralogs per enzyme. CYP4G and CPR are both single copy genes. **c:** HCR labelling of *CYP4G* transcripts (green) in oenocytes of the *Platyusa* abdomen (segments A4-A7 shown). Magenta: Hoechst-labelled nuclei. **d:** HCR labelling of *Platyusa CYP4G* (green), *FAS3* (magenta), *FAR4* (blue), nuclei (white) **e:** CYP4G-RNAi in adult *Platyusa* depletes total CHCs; p=.0.0017. **f:** δC13 values of specific CHCs from *Sceptobius* and *Liometopum* measured via GC-IRMS. **g:** Behavior arenas for tracking ant-beetle interactions. **h**: *Platyusa* demonstrating stereotyped ant appeasement behavior in arena. **i**: *Sceptobius* exhibiting stereotyped ant grooming behavior in arena. **j:** *Platyusa* appeasing *Liometopum* using substance secreted from abdomen tip. **k**: Contact time between beetles and *Liometopum* in behavioral arenas over 24 h. Asterisks denote p<0.0001 in Tukey post-hoc tests. **l:** *Sceptobius* climbing onto and grooming *Liometopum*. **m:** *Sceptobius* grasps the ant’s antennal scape with its mandibles to permit grooming. **n:** Baseline-corrected deuterated hydrocarbon transferred from *Liometopum* to beetles in behavioral arena over 24 h. Circled data points are cases of *Platyusa* attacking and consuming worker ants. Asterisks denote p<0.0001 in Tukey post-hoc tests. **o:** *Sceptobius* tarsi bear dense “grooming setae”, absent in free-living outgroup beetles in the tribe Falagriini.

We further tested this by isolating *Platyusa* from ants, and found the beetle can complete its life cycle in the laboratory in the absence of its host, reared on a diet of freshly killed *Drosophila*. Under these conditions, adult *Platyusa* present a CHC profile closely matching that of wild-caught beetles from *Liometopum* nests, albeit with altered ratios of certain compounds, implying beetles endogenously synthesize CHCs, and a shifted dietary input mildly influences the profile (**Fig. S2A**). We confirmed this by inhibiting endogenous CHC biosynthesis in *Platyusa*. Insect CHCs derive from a conserved oenocyte pathway in which long-chain fatty acids are elongated and reduced to aldehydes before decarbonylation, yielding secreted, very-long-chain alkanes and, if additionally desaturated, alkenes^73^. Using RNA-seq and differential expression analysis we recovered an entire CHC pathway expressed in abdominal tissue, with multiple paralogs encoding enzymes for fatty acid synthesis (FASN), elongation (ELO), reduction (FAR) and desaturation (desat) (**Fig. 2b, Figure S3, Table S2**). Some of these are orthologs of enzymes previously shown to mediate CHC biosynthesis in the free-living aleocharine, *Dalotia coriaria*^64^ (**Table S3, File S1**). Importantly, the terminal decarbonylation step in CHC biosynthesis is performed by a single, insect-specific cytochrome P450 enzyme, CYP4G^87^. We confirmed that *Platyusa CYP4G* is expressed in large abdominal cells intermingled with fat body tissue (**Fig. 2c**), which co-express copies of the other pathway enzymes (**Fig. 2d, Fig S4a**). Based on gene expression and anatomical location, we infer these cells to be oenocytes. In *Dalotia*, RNAi silencing of *CYP4G* causes loss of CHCs on the beetle’s body surface^64^, and injecting double-stranded RNAi into wild-caught *Platyusa* similarly reduced the levels of all CHCs (**Fig. 2e, Fig S5**). We conclude *Platyusa* synthesizes an approximate mimetic profile *de novo*, which imperfectly matches the CHC template of any specific host colony (**Fig. 1b, e-h**).

## Accurate chemical mimicry through horizontal CHC transfer

The mode of CHC mimicry in *Sceptobius* is strikingly different to that of *Platyusa*. We were able to resolve five CHCs suitable for compound specific GC-IRMS in *Sceptobius* beetles and ants from the same colony. In contrast to *Platyusa*, the δ^13^C values of each *Sceptobius* compound showed no significant offset from the corresponding ant compound, indicating a common biosynthetic origin (**Fig. 2f**). Within a nest, *Liometopum* workers outnumber *Sceptobius* beetles by 2-3 orders of magnitude, implying that the source of *Sceptobius* CHCs is likely the ant. We explored the basis of the differing modes of CHC mimicry employed by *Sceptobius* and *Platyusa* by tracking individual beetles and ants in behavioral arenas for 24 h (**Fig. 2g**). *Platyusa* avoided ants, spending <5% of the time in ant contact, similar to free-living *Dalotia* rove beetles (**Fig. 2k**). During brief encounters we observed *Platyusa* evade scrutiny by ants by “appeasing” them with secretions from the abdominal tip (**Fig. 2h, j**). In contrast, *Sceptobius* spent >30% of the time in prolonged, physical contact with *Liometopum* (**Fig. 2k**). Closer examination revealed a remarkable behavior in which *Sceptobius* mounts the dorsal ant body (**Fig. 2l**), clasping the ant’s antennae in its mandibles (**Fig. 2m**). Anchored to the ant in this way, the beetle grooms the ant by scraping its tarsi repeatedly over the surface of the ant, and then the surface of its own body (**Video S2**). We established that this behavior transfers CHCs from ant to beetle. Applying deuterated hydrocarbons onto *Liometopum* workers, and then housing these ants with beetles, we measured significant transfer onto *Sceptobius* after 24 h (**Fig. 2n, Fig. S6**), but negligible transfer onto *Platyusa* or *Dalotia*. *Sceptobius* tarsi bear specialized “grooming setae” that likely enhance CHC transfer via increased surface area (**Fig. 2o**). We conclude that *Sceptobius* horizontally “steals” CHCs from ants, achieving perfect mimicry of any gestalt, colony-specific CHC template (**Fig. 1c, e-h**).

## CHC production by *Sceptobius*

Our findings show that CHCs on the *Sceptobius* body originate from ants. Curiously, however, on assembly of a genome and whole-body transcriptome of the beetle we discovered an entire CHC pathway comprising intact (non-pseudogenized) loci for each catalytic enzyme. Multiple paralogs encoding FASs, ELOs, FARs and desats are present, as well as CYP4G (**Fig. 3a, Fig S7, Table S4**). Moreover, these enzymes are co-expressed in abdominal cells that we infer to be oenocytes (**Fig. 3b, c, Fig S4b**). This counterintuitive finding led us to consider what function CHCs may play in this species given its tight integration inside colonies, where it horizontally acquires the colony template. We speculated that the pleiotropic, non-pheromonal function of CHCs in desiccation resistance may be important (**Fig. 3d**). Over the years of this study, we conducted natural history observations, piecing together the life cycle of *Sceptobius* (**Fig. 3e**). We found that adult females produce giant eggs that fill the entire abdomen, (**Fig. S8**) and contain sufficient resources to support development from egg to adult. Females oviposit in damp soil directly at nest entrances; larvae hatching from these eggs do not need to feed, developing and pupating in soil, insulated from ants walking above them. After roughly two weeks, teneral (depigmented) adult beetles eclose, and search for an ant to groom; due to the proximity of egg laying to *Liometopum* nests, the groomed ant will likely belong to the beetle’s maternal colony, into which the beetle will integrate after “stealing” the CHC template (**Fig. 3e**).

**Figure 3.**
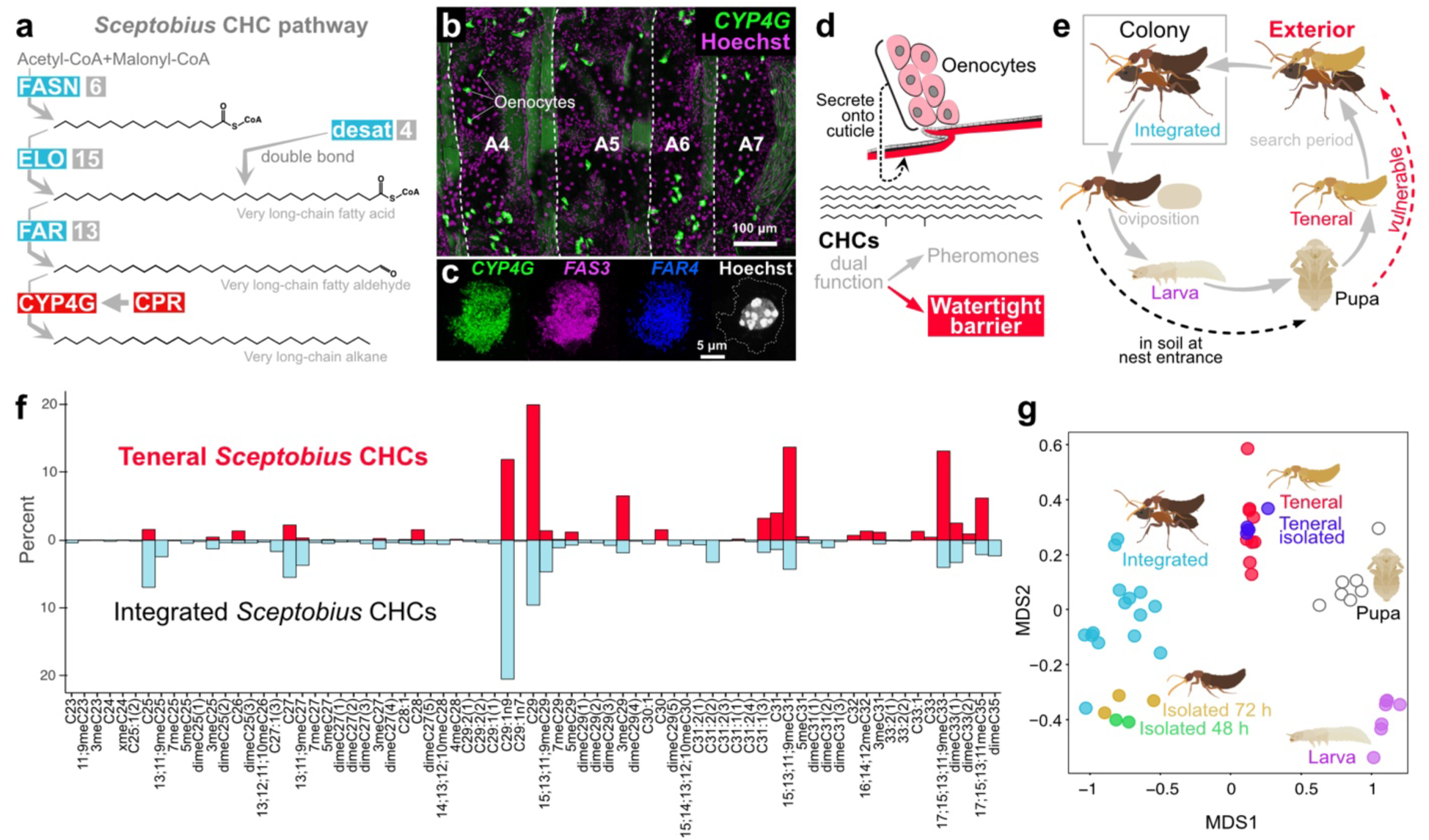
*Sceptobius* CHC biosynthesis. **a**. *Sceptobius* CHC pathway, with paralog numbers in grey boxes. **b:** *CYP4G* HCR (green) reveals oenocytes distributed in abdominal segments A4-A7 (magenta: nuclei). **c.** HCR reveals co-expression of CHC pathway enzymes in *Sceptobius* oenocytes (green: *CYP4G*; magenta: *FAS3*; blue: *FAR4*; white: nuclei). **d:** CHCs play a dual role as contact pheromones and a desiccation barrier on the cuticle. **e:** *Sceptobius* life cycle is split between the nest interior and exterior. Oviposition as well as larval and pupal development occur outside of the nest. After eclosing, teneral beetles groom ants, integrating into the nest interior. **f:** Teneral *Sceptobius* produce a CHC profile distinct from integrated *Sceptobius,* and thus also different from the host ant. **g:** NMDS ordination of CHC profiles measured across *Sceptobius* development, using Bray Curtis dissimilarity, 2D stress=0.05.

Prior to grooming, the teneral beetle is highly vulnerable to desiccation, bearing no ant-derived CHCs on its body (**Fig. 3e**). We collected *Sceptobius* eggs and larvae from *Liometopum* nest entrance soil and transferred them to ant-free soil to undergo metamorphosis. Remarkably, teneral beetles, pre-ant contact, bear a detectable CHC profile that must be produced endogenously, and is consistent across teneral beetles from different colonies (**Fig. 3f, g**). The teneral profile is clearly distinct from that of a mature, integrated beetle that has obtained its CHCs via grooming (**Fig. 3f, g**). The teneral profile instead appears to be a “skeletal” mimetic profile, comprising only 31 components, almost all of which are present in *Liometopum* (**Fig. 3f**). The profile lacks the full, 73-component complexity of a horizontally acquired CHC profile present on mature, integrated beetles (**Fig. 3f**). CHCs are also produced by *Sceptobius* larvae and pupae— each stage synthesizing a distinct profile (**Fig. 3g**). Hence, early *Sceptobius* life stages that inhabit the nest exterior produce CHCs, in keeping with the essential role of these compounds in desiccation resistance.

## *Sceptobius* develops a stealth phenotype via CHC pathway silencing

The total mass of endogenous CHCs on teneral *Sceptobius* is similar to that on *Sceptobius* larvae and pupae, and an order of magnitude less than the CHC-mass acquired by integrated beetles via grooming (**Fig. 4a**). Nevertheless, the incongruence between the teneral CHC profile and the mimetic profile obtained via grooming is of sufficient magnitude that were these endogenous CHCs still present on the body of an integrated beetle, they would disrupt perfect mimicry (**Fig. 4b**). NMDS ordination of the integrated *Sceptobius* CHC profile shifted, *in silico*, by the adding the average teneral beetle profile to it, would push most of the integrated profiles outside of ant CHC space (**Fig. 4c**.) How, then, does *Sceptobius* achieve perfect mimetic accuracy through grooming, when it is simultaneously capable of synthesizing a mimicry-disrupting, endogenous profile? We noted that, while imaging gene expression in oenocytes of mature, integrated beetles, transcript levels of CHC pathway enzymes, such as *CYP4G*, were barely detectable (**Fig. 4d**). We quantified CYP4G expression over the time course of early adult life, and found that expression is maximally high on eclosion from the pupa, dropping to approximately 30% by 72 h, before strongly diminishing to the baseline seen in integrated beetles (**Fig. 4d, e; Fig. S9a-c**). A parallel drop is observed for other CHC pathway enzymes (**Fig. S9, Fig. S10**). We infer that *Sceptobius* produces CHCs during the vulnerable pupal and teneral life-stages, which likely helps to counter desiccation (and may also be essential for successful eclosion from the pupal exuvium^88^). The teneral profile is minimal in amount and quasi-mimetic. During the subsequent search phase, in which the beetle locates a worker ant to groom, the CHC pathway undergoes transcriptional silencing, creating a chemically “stealth” beetle phenotype that can now achieve perfect chemical mimicry via grooming (**Fig. 4f**).

**Figure 4.**
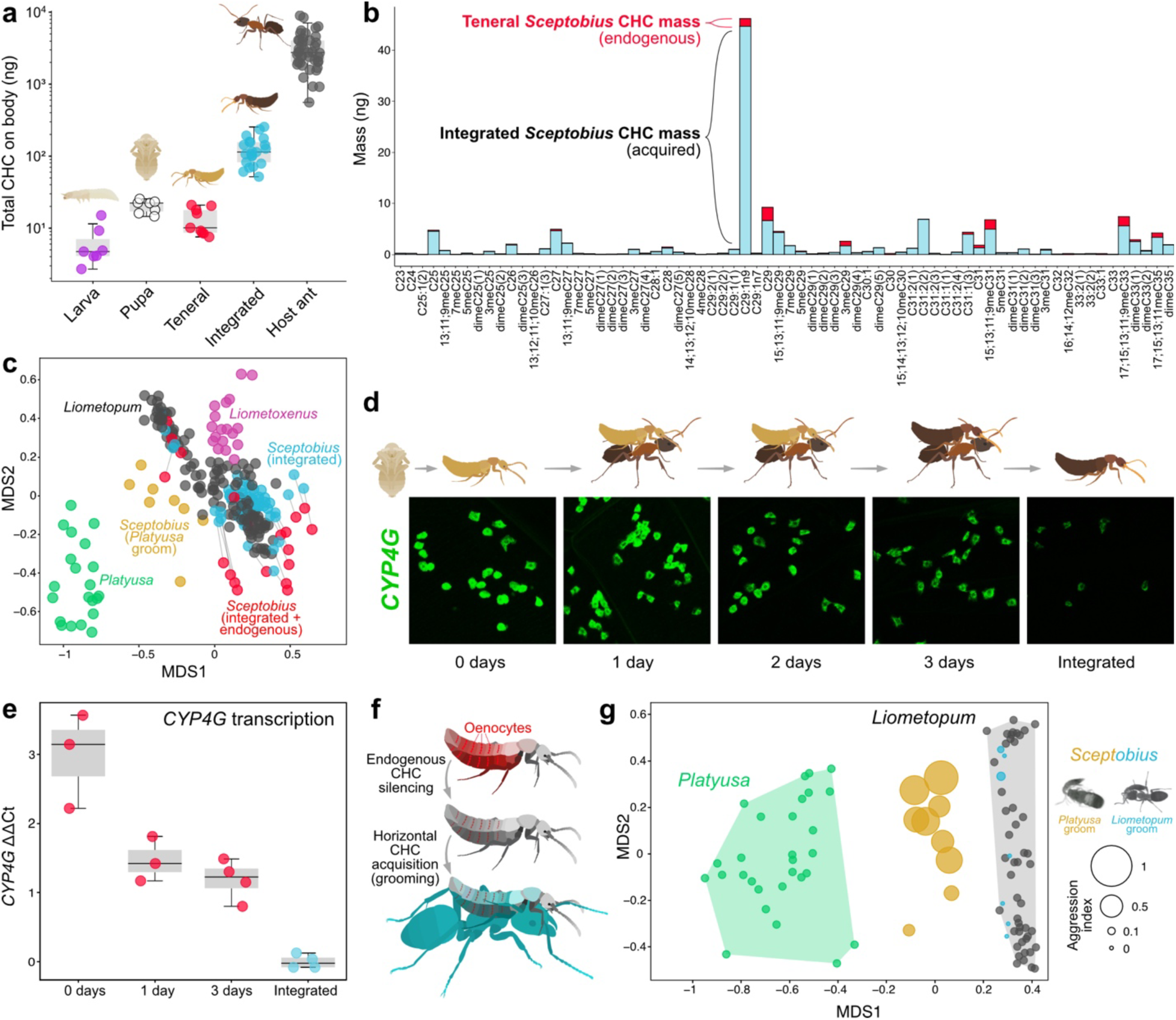
Adult *Sceptobius* develop a stealth phenotype. **a:** Total CHC mass on *Sceptobius* life stages and *Liometopum* workers. **b:** Mean amount of each CHC on integrated *Sceptobius* body (light blue), with the mean teneral profile stacked on top (red). **c:** NMDS ordination of *Liometopum* and myrmecophile CHC profiles (Bray Curtis dissimilarity, 2D stress=0.12). Blue data points are integrated *Sceptobius*; red points are combined integrated+teneral profiles (if integrated *Sceptobius* synthesized the teneral CHC profile in addition to acquiring CHCs from *Liometopum*). Gold points are profiles of *Sceptobius* that have groomed *Platyusa*. **d:** HCR timecourse of *CYP4G* expression levels in *Sceptobius* oenocytes, from eclosion from pupa to social integration in nest. **e**: qPCR timecourse of whole body *CYP4G* transcripts post-eclosion. **f:** *Sceptobius* silences CHC biosynthesis, horizontally obtains *Liometopum* CHCs via grooming to integrate into colonies. **g**: NMDS ordination of *Platyusa*, *Sceptobius*, and *Liometopum* CHC profiles (Bray-Curtis dissimilarity, 2D stress=0.08). CHC profiles for *Sceptobius* (with additional *Platyusa* CHCs: gold; control beetles: blue). Data point size indicates degree of aggression experienced by each beetle in a behavioral trial with three ants from the beetle’s colony of origin.

We performed an experiment which demonstrates that the silencing of endogenous CHC biosynthesis is essential for the social integration of *Sceptobius* into host ant colonies. In a separate study, we have found that *Sceptobius* recognizes its *Liometopum* host by detecting the ant’s CHCs, which elicit the beetle’s grooming behavior^89^. Because *Platyusa* presents the same CHCs as the ant, *Sceptobius* will similarly groom this other myrmecophile species^89^. We repeated this experiment and found that grooming *Platyusa*’s “approximately” mimetic profile pulls formerly integrated *Sceptobius* beetles outside of ant chemical space—a near-equivalent degree to the hypothetical addition of the teneral profile (**Fig. 4c**). Remarkably, disruption of the beetle’s CHC profile in this way breaks the stealth beetle’s cover: such beetles are attacked repeatedly by the ant, indicating a high fitness cost to endogenous CHC production inside the nest (**Fig. 4g**).

## Horizontal acquisition of ant CHCs is essential for *Sceptobius* survival

Our findings reveal how *Sceptobius* transitioned from a free-living beetle to a myrmecophile via two phenotypic innovations: i) a mechanism to silence endogenous CHC production, and ii) a social behavioral program to obtain CHCs via ant grooming. Through these twin modifications, the beetle gained the ability to integrate into colonies. This has come at the expense of being able to live away from ants. In contrast to *Platyusa*, isolating *Sceptobius* from ants leads to rapid mortality over 12–72 h, while under the same laboratory conditions the beetles survive long-term if ants are present (**Fig. 5a**). Just prior to and immediately following death, isolated beetles exhibit negligible levels of CHCs on their bodies due to rapid turnover of the previously acquired profile (**Fig. 5b**). The remnant profiles are close to those of integrated beetles, and clearly distinct from teneral beetles (**Fig. 3g**). We infer that silencing of endogenous CHC production is irreversible: isolated *Sceptobius* is incapable of compensatory biosynthesis of these compounds. This inability to reactivate biosynthesis implies that beetles are likely apart from ants at most transiently during adult life once integrated, removing selection pressure to endogenously produce these compounds.

**Figure 5.**
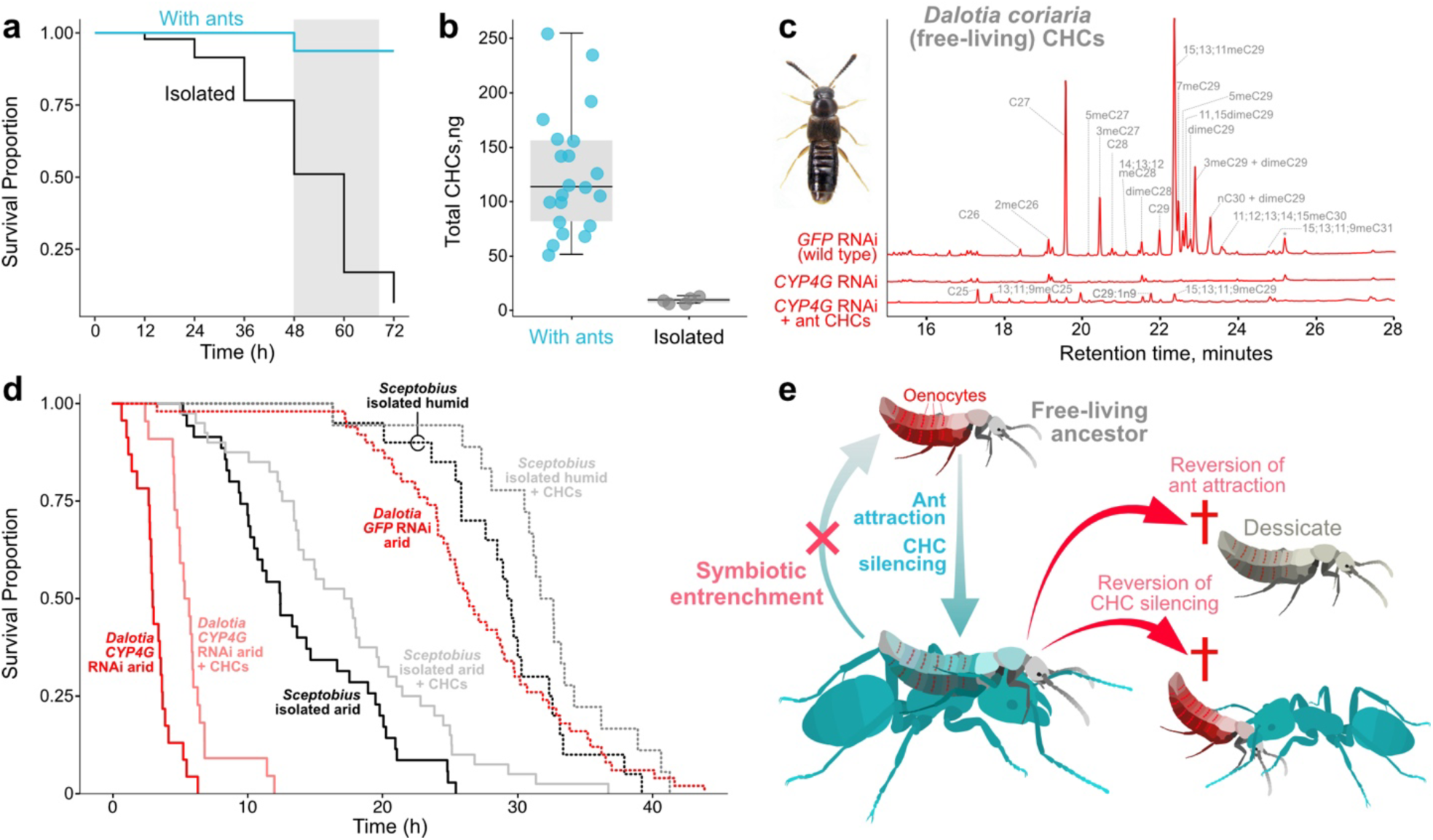
Entrenchment of myrmecophily. **a:** *Sceptobius* undergoes rapid mortality when physically isolated from ants. **b.** Isolating *Sceptobius* results in a drastic reduction of CHCs prior to death. **c:** Free-living *Dalotia coriaria* CHC production can be silenced, and with *Liometopum* CHCs. **d**. Survival curves of *Sceptobius* and *Dalotia* under CHC or environmental manipulations. **e**: Entrenchment of obligate myrmecophily in *Sceptobius* through reciprocal sign epistasis between ant attraction and CHC silencing.

We explored whether irreversible CHC loss may contribute to obligate dependence of *Sceptobius* on ants, contrasting this myrmecophile with *Dalotia*—an aleocharine approximating the free-living phenotype from which *Sceptobius* evolved^65^. *Dalotia* endogenously produces CHCs, which can be removed by silencing *CYP4G* (**Fig. 5b**). In arid conditions, wild-type *Dalotia* survival begins dropping at ∼20 h, reaching 50% at ∼28 h (**Fig. 5c**). Silencing *CYP4G* renders *Dalotia* desiccation-prone, the beetles now reaching 50% mortality within ∼5 h (**Fig. 5c**). Application of a small level of *Liometopum* CHCs to such *Dalotia* (**Fig. 5c**) moderately but significantly prolongs survival (**Fig. 5d**) These results underscore the critical function that CHCs play in water balance and survival of free-living beetles. We permitted *Sceptobius* to groom ants and acquire a full CHC profile before isolating beetles in the same arid conditions used for *Dalotia*. Such beetles begin dying rapidly, reaching 50% mortality within 12 h, implying that ant CHCs are quickly lost (**Fig 5d**). Repeating this experiment with application of additional *Liometopum* CHCs moderately prolonged *Sceptobius* survival, while increasing humidity approximately doubled lifespan—*Sceptobius* survival now approximating that of free-living *Dalotia* under arid conditions (**Fig. 5d**). Some water loss through the chitinous cuticle can still occur in humid condtions^90^, hence even at elevated humidity the application of additional *Liometopum* CHCs partially extended *Sceptobius* survival (**Fig. 5d**). CHCs acquired from ants are therefore critical for *Sceptobius* survival. We infer that the evolution of grooming behavior not only achieved chemical mimicry and social integration; it compensated for the loss of endogenous CHCs that were essential for desiccation resistance in *Sceptobius*’ free-living ancestors.

## Discussion

The phenotype we report for *Sceptobius* represents a putative syndrome that has evolved convergently in aleocharine beetles. Myrmecophile lineages that are socially integrated are often those that groom worker ants (or are groomed by them)^60,91–97^ and undergo rapid mortality on isolation^91^ (**Fig. S11a-c**). All such clades show no instances of reversion to living freely^60^—a macroevolutionary pattern for which we offer a mechanistic explanation. We have shown that CHC silencing in adult *Sceptobius* has become an essential trait. Reversion of this phenotype to one of sustained CHC biosynthesis would be costly, breaking perfect mimicry in a beetle that has evolved to physically interact with ants (**Fig. 5e**). Concurrently, the beetle’s strong physical attraction to ants has also become essentialized. Its loss would be lethal, since the ant has become the source of CHCs that form a critical desiccation barrier (**Fig. 5e**). *Sceptobius* is thus caught in an ecological Catch-22 where loss of either symbiotic trait renders the other deleterious. This arrangement is likely to have a ratcheting effect on evolution—the low probability of simultaneous reversion locking the lineage into obligate ant dependence (**Fig. 5e**). Entrenchment may therefore be inherent to a form of social integration that has evolved convergently and irreversibly across this vast beetle subfamily. Accordingly, trait loss, such as the convergent degeneration of eyes, wings and the defensive tergal gland in these taxa (**Fig. S11d, e**), may be a later consequence of this primary entrenching mechanism.

We do not know the underlying genetic architecture of each symbiotic trait in *Sceptobius*, but the Catch-22 holds even if single mutational changes could eliminate either trait (or cause its reversion to the ancestral state). Reciprocal dependence between the two traits excludes their individual reversion. This condition, where two traits that permit symbiosis are functional when together but deleterious alone, is a phenotypic manifestation of “sign epistasis”, in which a mutation’s cost or benefit is contingent on the genetic background in which it occurs^98^. Sign epistasis underlies irreversibility at the protein level, where substitutions that alter protein function^99–101^ or subunit binding^102^ become entrenched through epistatic interactions with more recent substitutions. We propose that an analog underlies cases of symbiotic entrenchment. Traits that promote interactions with other organisms may undergo loss or reversion if they are epistatically unconnected or show non-reciprocal epistasis, offering potential return paths to living freely. In *Platyusa*, closely associating with ants is likely only beneficial in the context of its ability to endogenously mimic its host’s CHCs, but endogenous CHC mimicry is not itself deleterious in the absence of ants. Loss of ant attraction may therefore be possible in *Platyusa*, permitting reversion of myrmecophily in this species. In contrast, adaptive or neutral evolution of, at minimum, two distinct traits promoting symbiosis will foster entrenchment should mutational loss or reversion of one of these traits render a second one deleterious. Such is the scenario we demonstrate between ant attraction and acquired CHC mimicry in *Sceptobius*.

Reciprocal sign epistasis, operating in the transition to symbiosis, may be a general force that biases the direction of evolution towards greater integration with hosts. We note that many symbioses, including endosymbiosis^103,104^, ecto- and endoparasitism^105–107^, and social parasitism^81,108^, hinge fundamentally on mechanisms for finding and/or physically associating with hosts, together with body or cell surface modifications that achieve compatibility with host immune or sensory surveillance. Pleiotropy of an organism’s surface as both a barrier to the environment and the bearer of self/nonself information, positions it in potential reciprocal epistasis with host-interacting mechanisms. The Catch-22 we identify for *Sceptobius* may therefore be a common basis for symbiotic entrenchment across the tree of life.

*****

## Acknowledgements

We are grateful to Matt Pennell, Michael Dickinson and members of the Parker lab for critical feedback on this manuscript. This study was supported by funding to JP from the National Science Foundation (CAREER 2047472), the Army Research Office (MURI W911NF18S0003), an Alfred **A.** P. Sloan Fellowship, a Pew Biomedical Scholarship, Rita Allen Scholarship, Klingenstein-Simons Fellowship, and research Grants from the Shurl and Kay Curci Foundation and Okawa Foundation.

## Data Availability

Data were uploaded to CaltechData: https://data.caltech.edu/records/bw6tm-ey076

## Methods

### Specimen collection

Beetles and ants were collected from *Liometopum occidentale* nests in the Angeles National Forest between 2017 and 2024. *Liometopum* nests were most commonly located in Coast Live Oak (*Quercus agrifolia*) and California Bay Laurel (*Umbellularia californica*). Specimens of all three rove beetle species were primarily collected by sieving leaf litter and soil surrounding nests at the bases of trees. *Sceptobius* was also collected from internal nest material, while both *Sceptobius* and *Liometoxenus* were additionally collected from nest entrances. *Platyusa* eggs and larvae were collected in leaf litter surrounding *Liometopum* nests. *Sceptobius* eggs and larvae were collected in damp soil directly at nest entrances, comprising frass formed from the heartwood excavated by *Liometopum*. *Platyusa* and *Sceptobius* were collected year-round (peaking between February and August). *Liometoxenus* is seasonal and was only collected between February and May. Beetles and ants were aspirated and placed into 50 ml tubes filled with damp kimwipes, and transported back to the laboratory.

### Myrmecophile culture

In lab, beetles were maintained in large plastic bins (Rubbermaid) until they were used in experiments. *Platyusa* were reared without ants in containers with a layer of densely packed, damp coconut fiber. Frozen *Drosophila melanogaster* or *Liometopum occidentale* workers were placed in containers on pieces of foil as a food source. *Platyusa* reared in lab would periodically lay viable eggs that would develop into adults. *Platyusa* eggs and larvae were additionally collected in leaf litter at the field sites, or in leaf litter brought back to the lab and sorted under a microscope. They were maintained in lab on damp coconut fiber substrate in the same containers as adult *Platyusa*. Larvae were fed frozen *Drosophila* and proceeded through three larval instars prior to pupation. Both *Sceptobius* and *Liometoxenus* were kept in containers with *Liometopum* workers collected from the same nest. The walls of these containers were coated in InsectaSlip/Fluon (Bioquip, CA) to prevent ant escapes. Ants were provided with sucrose solution and glass tubes partially filled with water and cotton ball stoppers to provide moisture. Soil was brought back to lab where *Sceptobius* eggs and larvae were located in soil using a stereomicroscope. Eggs were stored on slightly damp filter paper in petri dishes and observed daily for the presence of freshly hatched larvae. Larvae were transferred to fresh petri dishes with damp filter paper. Attempts to identify a food source for *Sceptobius* larvae were unsuccessful and was found that *Sceptobius* larvae do not require feeding to successfully pupate. Prior to pupation, the larvae pass through two larval instars. The container of pupae was checked twice daily for the presence of freshly eclosed teneral *Sceptobius.* Teneral *Sceptobius* died rapidly if not introduced to *Liometopum* within a day of eclosing. Teneral *Sceptobius* not being sampled on the day of eclosion were placed in petri dishes with damp filter paper and five *Liometopum* workers. The ants were kept alive by placing sucrose solution-soaked filter paper squares on pieces of foil in petri dishes.

### Cuticular Hydrocarbon analysis

Insects were extracted in 70 µL hexane (n-hexane SupraSolv®, Sigma Aldrich, MO) for 20 minutes. Samples for which semi-quantification of CHC amounts was required were extracted in hexane containing a 10 nanogram-per-microliter octadecane (C18, Sigma Aldrich, MO) internal standard. Crude extracts (2µL) were then injected into a Shimadzu QP2020 GCMS with an AOC-20i Shimadzu autosampler. The instrument was equipped with Helium as a carrier gas and a Phenomenex (Torrance, CA) ZB-5MS fused silica capillary column (30m x 0.25 mm ID, df=0.25µm). The injection port was operated at 310°C in splitless mode, with a column flow rate of 2.15 mL/min. The column was held at 40°C for 1 minute, ramped at 20°C/min to 250°C, ramped at 5°C/min up to 320°C, and then held at 320°C for 7.5 minutes. The transfer line was held at 320°C and the ion source temperature was held at 230°C. Electron ionization was carried out at an ion source voltage of 70eV, and MS scans were collected between 40 m/z and 650 m/z at a scan rate of 2 scans per second.

For some organisms, including *Oligota* and some of the *Sceptobius* samples, beetles were extracted in 10 microliters of hexane due to the low amount of CHCs on the organisms. When quantification of these samples was required, hexane with a 1 ng/microliter C18 internal standard was used. Low volume samples were manually injected into the GCMS. Identification of individual CHC compounds was determined based on retention index as well as diagnostic ions, and comparison to previously described *Liometopum* and *Drosophila* GCMS data^109,110^ GC peaks were manually integrated using LabSolutions Postrun Analysis (Shimadzu, Kyōto, Japan). Alkene double bond position was previously identified for a number of *Liometopum* compounds^110^ Absolute CHC amounts were calculated by taking the ratio of the area of each peak to the C18 internal standard peak area, and multiplying by the total mass of internal standard in the extraction. CHC measurements were converted to percent composition for each spectrum. Pairwise Bray-Curtis dissimilarity was calculated across all samples and then non-metric multidimension scaling (NMDS) ordination was performed using the metaMDS function from the vegan R package^111^ The R function hclust was used to perform hierarchical clustering of the pairwise Bray-Curtis dissimilarity matrix. Hierarchical clustering was additionally performed on the pairwise Jaccard dissimilarity matrix of the binarized CHC data.

### Gas Chromatography Isotope Ratio Mass Spectrometry (GCIRMS)

Due to the high concentrations required for accurate measurement, ants and beetles were pooled with conspecifics from the same nest/collecting trip for stable isotope measurements. Between 28-50 ants, 9-64 Sceptobius, and 20 Platyusa were pooled for each extraction. Insects were extracted in 1mL of hexane, which was then evaporated under a stream of nitrogen to a volume of roughly 100 microliters. Samples were then analyzed on a Thermo Trace GC_ultra_ (Thermo Fisher, CA) interfaced to a Thermo-Scientific Delta_+_XP GC-combustion-IRMS (Thermo Fisher, CA) equipped with a ZB-5MS column (30m x 0.25 mm ID, df=1µm, Phenomenex). Samples were injected in splitless mode into a PTV injection port with He as the carrier gas. The column was initially held at a temperature of 80°C for 1 minute, followed by a 20°C/min ramp up to 250°C and then a 3°C/min ramp up to 320°C, which was held for 12 minutes. A CO_2_ reference gas (δ^13^C = - 32.4‰) was co-injected during each sample run, generating four reference peaks against which sample δ^13^C was calculated relative to VPDB (Vienna Pee Dee Belemnite). An external standard containing ethyl icosanoate (from the ‘F8 mix’ of Arndt Schimmelmann, Indiana University), which has a known δ^13^C value of -26.1‰, was run every eight samples. The difference between the measured value and true value of the ethyl icosanoate standards bounding each set of eight runs was used to correct the δ^13^C values of the intervening samples. Samples were measured in triplicate and arithmetic means are reported. Peaks were identified by comparison to GCMS measurements and the stereotypical elution order of the CHCs of the three organisms.

### Life stage analysis

First and second instar larvae, as well as prepupal larvae were extracted in 10 microliters of hexane containing 1 ng/microliter C18 for 20 minutes. The extracts were then manually injected (3µL) into the GCMS and run with the CHC-GCMS program described above. Neither the first, nor the second instar larvae possessed measurable CHCs. To obtain pupae for CHC analysis, prepupal larvae were left on damp filter paper in petri dishes at room temperature. Pupae were collected within one day of pupation and extracted and run on the GCMS in the same manner as the *Sceptobius* larvae. Teneral *Sceptobius* were generated by allowing larvae to pupate in soil, which had been cleared of *Liometopum*. As individual teneral beetles eclosed, they were collected and either immediately extracted for CHCs, placed into petri dishes with damp filter paper for isolation analysis, or placed into petri dishes with damp filter paper and multiple *Liometopum*, for later time point measurements. Isolated teneral *Sceptobius* were kept on damp filter paper in Petri dishes for 24 hours, after which they were extracted and run on the GCMS using the same method as the *Sceptobius* larvae and pupae. High mortality was observed for the teneral beetles during this 24 h isolation period.

### Deuterium transfer

Approximately 0.5 milligrams of fully deuterated tetracosane and triacontane (triacontane-d_62_ and tetracosane-d_50_, 98 atom % D, Sigma-Aldrich, MO) were added to a 5 mL glass vial along with 1 mL of hexane to fully dissolve the hydrocarbons. The hexane was then evaporated off under a steady stream of nitrogen gas leaving a thin film of the two deuterated hydrocarbons on the inner wall of the vial. *Liometopum* were added to the vial and then shaken for ∼1 minute to transfer the labeled hydrocarbons to the surface of the ants. Uncoated control ants were prepared in a similar fashion, using a vial treated only with hexane. Deuterated hydrocarbon labelled and control *Liometopum* were paired with either *Platyusa, Sceptobius,* or *Dalotia* in wells within an IR transparent black acrylic arena, **Figure 2G**. Interactions were recorded in the dark at 1 hz under IR illumination using a FLIR camera (BFS-U3-51S5M-C: 5.0 MP) with a Pentax 12mm 1:1.2 TV lens (Ricoh, FL-HC1212B-VG). After 24 hours, the arena was removed and ants and beetles were extracted in 70 microliters of hexane with 25ng/microliter of octadecane as an internal standard. Samples were run on Shimadzu QP2020 GCMS equipped with helium as a carrier gas and a Phenomenex ZB-5MS fused silica capillary column (30m x 0.25 mm ID, df=0.25µm). The injection port was operated at 310°C in splitless mode, with a column flow rate of 2.15 mL/min. The column oven was held at 40°C for 1 minute, followed by a 40°C/min ramp up to 250°C, a 20°C/min ramp up to 320°C, and then a 5 minute hold at 320°C. The transfer line was held at 320°C and the ion source temperature was held at 230°C. The MS was operated at an ion source voltage of 70eV and scans were collected in selected ion monitoring mode (SIM), monitoring m/z 66, 85, 98, 254, 389, and 485 at 3.33 scans per second. Videos were reviewed and only interactions in which the beetles had not been killed were used.

The C18, triacontane-d_62_, and tetracosane-d_50_ peaks were manually integrated (sum of the selected ion counts), and the absolute amounts of the deuterated hydrocarbons were calculated by taking the ratio of the area of each peak to the C18 internal standard peak area, and multiplying by the total mass of internal standard in the extraction. The average background signal was calculated for the control ants and beetles and subtracted from the corresponding treatment groups. The background-corrected deuterated hydrocarbon mass on each beetle was then divided by the corresponding measurement from the paired ant to calculate the ratio of deuterated hydrocarbon transferred in each interaction well. Differences in the transfer ratio between the three beetles were compared using an ANOVA test with a Tukey *post hoc* test in R.

### Phylogenomics

For phylogenomic inference, transcriptomes were first generated for *Sceptobius, Platyusa,* and *Drusilla*. These three transcriptomes were then combined with gene predictions from 26 other beetles and *Drosophila melanogaster*. Orthologous sequences were identified from the 30 genomes samples and a maximum likelihood species tree was calculated from a supermatrix constructed from single-copy gene families. Both *Platyusa sonomae* and *Sceptobius lativentris* were collected at Chaney Canyon, Angeles National Forest, California (34°13’00.9”N 118°09’09.9”W) during the summer of 2018-2019. *Drusilla canaliculate* specimens were collected by Tim Struyve (locality: Moerbeke, Belgium, 51°10’37.2”N 3°54’32.4”E, 04 iv 2017). Roughly 120 *Sceptobius* were dissected, separating the 6^th^ and 7^th^ abdominal segments from the 8^th^ abdominal segment, as well as legs and antennae from the rest of the body. Legs and antennae were further separated by sex and combined with ∼20 male and female *Sceptobius,* which were separated into pools of legs, antennae, and the rest of the body, separated by sex. One each of male and female *Platyusa* and *Drusilla* were separated into heads and bodies. Total RNA was extracted from live or flash frozen materials stored at -80°C using a ZYMO *Quick-RNA* Tissue/Insect extraction kit (ZYMO Research, CA). RNA quantity was assessed with a Nanodrop (Thermo Fisher, CA). *Sceptobius* 150bp libraries were prepared and sequenced paired end to a read depth of 100 million reads and *Platyusa* and *Drusilla* 100bp libraries were prepared and sequenced paired end to read depth of 100 million reads, all by Omega bioservices. The *Platyusa* and *Drusilla* reads were initially filtered by mapping raw reads from both species onto a concatenated *Platyusa*/*Drusilla* genome (generated previously^65^) using BOWTIE2^112^ due to account for any cross contamination of the samples during library prep. Only reads mapping to the correct portion of the combined genomes were used in subsequent analysis. Similarly, *Sceptobius* reads were filtered by mapping raw reads from all libraries onto a concatenated *Sceptobius/Dalotia coriaria* genome and only reads mapping to the correct species were used in downstream analysis. The filtered reads from *Platyusa, Drusilla,* and *Sceptobius* were then filtered using rCorrector to remove erroneous k-mers and filtered using a script located at: https://github.com/harvardinformatics/TranscriptomeAssemblyTools to remove uncorrectable reads. TrimGalore v0.6.0^113^ was used to remove any remaining adaptors from the reads. The *Platyusa, Drusilla*, and *Sceptobius* libraries were then fed to TRINITY v2.12.0^114^ for de Novo transcriptome assembly with Jaccard clipping. Assembled transcriptomes were assessed for completeness via gVolante^115^ using BUSCO v5^116^ with the Arthropoda orthologue set. BUSCO scores for *Platyusa*, *Drusilla*, and *Sceptobius* were 99.9%, 99.9%, and 100% respectively for the 1013 core genes analyzed. The *de novo* transcriptomes were then processed with transdecoder v5.5.0 to predict open reading frames and run through CDHIT v4.8.1^117^ using a 98% sequence identity threshold, -p 1 -d 0 -b 3 -T 10, to remove duplicate sequences.

The three transcriptomes were combined with gene predictions from 27 other genomes, including *Drosophila melanogaster,* various free-living beetles spanning the suborder Polyphaga, and the myrmecophile *Liometoxenus newtonarum*, all of which were collated or assembled previously^65^. Protein-coding sequences for the 30 species were processed with Orthofinder v2.5.2^118,119^ (-M msa -S diamond -A mafft -T fasttree) to find orthologous sequences. The resulting mafft amino acid sequence alignments for the orthogroups containing at least one sequence for at least 24 of the species were then trimmed with the trimal v1.4.1^120^ gappyout method, resulting in 2064 orthologous protein coding loci. Maximum likelihood trees were generated for all trimmed alignments using iqtree v1.6.8^121,122^ with 1000 bootstraps, restricted to the models WAG,LG,JTT, and Dayhoff. The trees were pruned to strictly orthologous sequences using PhyloTreePruner v1.2.4^123^ with a minimum number of taxa cutoff of 24 (80%) and a bootstrap cutoff of 0.7. The resulting pruned alignments were concatenated with FASconCAT v1.04^124^, resulting in a supermatrix containing 750,163 amino acid sites with 2,063 gene partitions. MARE v0.1.2^125^ was used to improve the phylogenetic signal of the partitions, resulting in a supermatrix containing 374,139 amino acid sites with 1,039 gene partitions. Partitionfinder v2.1.1^126^ was then used with raxml^127^ and otherwise default parameters to identify an optimal partitioning scheme and corresponding set of molecular evolution models for the gene partitions. A maximum likelihood species tree was generated from the supermatrix and partition scheme using iqtree v1.6.8 with 1000 bootstrap replicates.

In order to constrain divergence times on the species tree, twelve fossil calibrations were selected to place bounds on specific nodes in the tree. For all fossils, the youngest age interpretation of the fossil was used, following best practices^128^. The corresponding nodes are labeled in Figure S1a.

**A. MRCA of Diptera and Coleoptera:** An upper bound was placed on the root of the tree using the median age estimate of holometabola (345 Ma)^129^. A lower bound was set using the stem group beetle *Coleopsis archaica* (293.8 Ma)^130^.
**B. MRCA of Buprestoidea and all other beetles in this study:** A lower bound was placed on the node at which *A. planipennis* branches, using the stem Buprestid fossil, *Ancestrimorpha volgensis* (164.7Ma)^131,132^.
**C. MRCA of Curculionoidea and Chrysomeloidea:** A lower bound was placed on the MRCA of Curculionoidea and Chrysomeloidea using both *Archaeorrhynchus* sp. and *Eobelus* sp. (157.3Ma)^133,134^.
**D. MRCA of Chrysomeloidea:** A lower bound was placed on crown Chrysomeloidea using *Cretoprionus liutiaogouensis* (122.5 Ma)^134,135^.
**E. MRCA of Staphylinidae:** A lower bound was placed on the MRCA of Staphylinidae using undescribed Silphids (152 Ma)^136,137^.
**F. MRCA of Aleocharinae and Tachyporinae:** A lower bound was placed on the MRCA of Aleocharinae and Tachyporinae using the Tachyporine beetle *Protachinus minor* (147.28 Ma)^138,139^.
**G. MRCA of Gymnusini:** A lower bound was placed on the Gymnusini MRCA using *Cretodeinopsis aenigmatica* (98.17 Ma)^140–142^.
**H. MRCA of *Adinopsis* and *Deinopsis*:** A lower bound was placed on the MRCA of *Adinopsis* and *Deinopsis* using *Adinopsis groehni* (43.1 Ma)^143^.
**I. MRCA of Hypocyphtini:** A lower bound was placed on the MRCA of the hypocyphtines using *Baltioligota electrica* (43.1 Ma)^144,145^.
**J. Aleocharini stem:** A lower bound was placed on the *Aleochara* stem based on *Aleochara baltica* (43.1 Ma)^146^.
**K. MRCA of Oxypodini and Homalotini:** A lower bound was placed on the MRCA of Homalotini and Oxypodini using *Phymatura electrica* (43.1 Ma)^146^.
**L. MRCA of Athetini and Geostibini:** A lower bound was placed on the MRCA of Athetini and Geostibini using *Atheta jantarica. Dalotia coriaria* is most likely a member of the *Atheta,* thus the calibration point was moved one node deeper from the Athetini MRCA (43.1 Ma)^145^.

MCMCtree and codeml, from PAML v4.9^147^, were used to perform divergence time analysis, using the previously generated maximum likelihood species tree, the associated supermatrix, and the above fossil calibration points. Branch lengths were initially approximated in codeml by maximum likelihood (ML) using the WAG empirical rate matrix with gamma rates among sites, approximated with four rate categories. The gradient and Hessian of the likelihood function at the ML branch length estimates were then used to run MCMCtree with the approximate method. The relevant model parameters were clock = 2, cleandata = 0, BDparas = 1 1 0.1, rgene_gamma = 2 20 1, sigma2_gamma = 1 10 1, and finetune = 1:.1 .1 .1 .1 .01 .05. All fossil calibration points were modelled as truncated Cauchy distributions^148^, defined with default MCMCtree parameters, with the exception of the root, which was defined as a uniform distribution between the upper and lower bound with tails on either side of the bounds^149^. After a burn in of 20,000 iterations, 200,000 samples were collected, sampling every 100 iterations. Divergence time estimates were compared between multiple runs to verify that the MCMC chains had converged to a stable posterior distribution.

### SMART-seq transcriptome sequencing

The abdominal fat body and entire crushed pronotum were dissected from individual *Platyusa,* and the 5th-7th abdominal segments, with gut removed, and the entire crushed pronotum were dissected from *Sceptobius* in ice-cold DEPC PBS, flash frozen in a dry ice/ethanol bath, and stored at -80°C until processing. Library preparation was carried out using the NEBNext Single Cell/Low Input RNA Library Prep Kit for Illumina, using NEBNext Multiplex Oligos (New England Biolab, MA) following the manufacturers protocol (NEB #E6420). The number of PCR cycles during cDNA amplification was 9 for *Sceptobius* samples and 14 for *Platyusa* samples. Final library amplification was either 8 or 12 cycles for all libraries, depending on the intermediate library concentration at this step. The quality of all libraries was assessed by running samples on a Qubit High Sensitivity dsDNA kit (Thermo Fisher, CA) and Agilent Bioanalyzer High Sensitivity DNA assay (Agilent, CA). The libraries were sequenced, either 50bp for *Platyusa* or 100 bp for *Sceptobius*, single end to a read depth of 20-25 million reads on Illumina HiSeq2500 (Illumina, CA) at the Millard and Muriel Jacobs Genetics and Genomics Laboratory at Caltech.

### Differential Expression Analysis

SMART-seq reads were pseudoaligned to transcriptomes of *Sceptobius* and *Platyusa* using kallisto v0.46.2, with 100 bootstraps, single end flag, an average fragment length of between 339-450bp with a standard deviation of 27-41bp. Two paired *Sceptobius* samples from the same animal were removed after mapping, due to the small percentage of reads that pseudoaligned (<4% vs 20-40% for all other samples). Next, sleuth v0.30.0 was run, using a full model of condition (pronotum or fat body) plus animal (paired samples were collected from each beetle) and a reduced model of animal. A likelihood ratio test was run using the two models to identify differentially expressed transcripts, which were then filtered by the sign of the average transcripts per million (TPM) for the transcript in the abdominal fat body minus the average TPM for the transcript in the pronotum.

### Identification of CHC pathway enzymes

All transcripts in the *Sceptobius* and *Platyusa* transcriptomes were searched against the NCBI nr (February 2019) and UniProt (February 2019) databases, taking the top five hits and using an evalue cutoff of 1e-05. The top hit for each transcript ID, sorted by highest bitscore, lowest evalue, and highest percent identify, was determined for both the NCBI nr and Uniprot results. The differentially expressed smartseq transcripts were then filtered by the NCBI nr and Uniprot annotations, selecting those transcripts annotated as elongation of very long chain fatty acids, fatty acyl-CoA reductase, fatty acid synthase, acyl-CoA desaturase, stearoyl-CoA desaturase, cytochrome P450 4G1/4G15, or NADPH-cytochrome P450 reductase. The orthogroups containing these potential CHC pathway enzymes were identified.

The enzyme list generated via differential expression analysis was supplemented with a homology based approach, drawing from cuticular hydrocarbon biosynthesis literature in *Drosophila melanogaster*^150,151^ and the closely related rove beetle, *Dalotia coriaria*^64^. Orthogroups containing known *Dalotia* and *Drosophila* CHC biosynthesis enzymes as well as the differentially expressed *Sceptobius* and *Platyusa* enzymes were analyzed by aligning the sequences in each orthogroup using mafft v7.453^152^ and then generating phylogenetic trees using fasttree v2.1.10^153^ (**File S1**). The resulting gene trees were manually curated, pruning analogous branches, removing short sequences that appeared to be fragmented, removing or splitting long sequences that appeared to be mis-predicted chimeras of adjacent genes in the genome, removing identical or nearly identical sequences, and in some cases replacing misassembled transcripts with gene predictions from the corresponding draft genomes. All *Sceptobius* and *Platyusa* sequences in each gene tree were assumed to be potential CHC biosynthesis enzymes. Curated transcriptomes for both *Sceptobius* and *Platyusa* were generated by replacing or removing those sequences modified or removed in the process of generating the gene trees. The smartseq reads were then remapped, in a similar fashion as described above, to these curated transcriptomes and differential expression analysis was performed. Heatmaps of TPM for the full complement of enzymes as well as their differentially expressed status in *Sceptobius* and *Platyusa* are shown in **Figures S4** and **S5.**

### Hybridization Chain Reaction (HCR)

Probe sets, HCR hairpins, as well as amplification buffer, hybridization buffer, wash buffer were purchased from Molecular Instruments (Beckman Institute at Caltech; www.moleculartechnologies.org) for each transcript. Representative transcripts were chosen from each gene family in *Sceptobius* and *Platyusa* using several criteria. As the only representative of its family, *CYP4G1* was chosen. *Desat1* is homologous to *Drosophila desatF*, *desat1*, *desat2*, and thus was chosen as a likely oenocyte enzyme candidate. *ELO4* was chosen due to its known expression and function in *Dalotia* oenocytes (the orthologous gene is named elo-708 in Brückner et al.^64^)*. FAS3* and *FAR4* were chosen due to their differential expression in the abdomen as well as minimal expression in the pronotum control tissue.

Samples were dissected in DEPC treated PBS with a small amount of DEPC PBST (0.1% Tween) added. Beetles were CO_2_ anesthetized and then transferred to a dissecting dish and the head was removed with dissecting scissors. Grasping the thorax and abdominal segments 9 and 10 with forceps, the abdominal tip, gut, and genitalia were removed through the tip of the abdomen. The abdomen was then either separated along the sagittal plane or ventrally and dorsally with dissecting scissors. Samples were placed in ice-cold DEPC-PBST until fixing. Abdominal segments were fixed in 4% PFA in DEPC-PBST for 25 minutes at room temperature and subsequently rinsed three times with fresh DEPC-PBST. Samples were then dehydrated by washing with a DEPC-PBST/methanol series, finishing with a 100% methanol wash. At this step, samples were either stored at -20°C or rehydrated using a methanol/DEPC-PBST series. Samples were rinsed two additional times for five minutes each at room temperature and then washed with a 0.01% Proteinase K solution in DEPC-PBST for five minutes at room temperature. Samples were then rinsed twice with fresh DEPC-PBST and post-fixed with a 4% PFA in DEPC-PBST solution for 25 minutes at room temperature. Samples were rinsed again with DEPC-PBST and then incubated with hybridization buffer at 37°C for 30 min. The probe solution was preheated during this period, combining two microliters of both even and odd HCR probes with 100 µL of hybridization buffer at 37°C. The hybridization buffer was removed from the sample and replaced with probe solution.

Samples were incubated overnight at 37°C. No-probe control samples were incubated overnight at 37°C in hybridization buffer in the absence of probes. The following day, samples were washed with preheated wash buffer, twice for five minutes and twice for 30 minutes, both at 37°C. During the last wash, hairpins were snap cooled (90 seconds at 95°C and then 30 min room temp in the dark). Samples were incubated with amplification buffer for 10 min, after which the old amplification buffer was replaced with 100 µL of fresh amplification buffer and the hairpins. Amplification occurred overnight in the dark at room temp. Multiple combinations of B1, B3, and B4 amplifiers with Alexa 488, Alexa 546, and Alexa 647 fluorophores were used. Samples were then washed with SSCT (saline sodium citrate with Tween) twice for five minutes in the dark. Samples were incubated for two hours at room temperature with Hoechst 33342 (1:2000, Thermo Fisher, CA) to mark nuclei, Alexa 488-Wheat Germ Agglutinin conjugate (WGA; 1:500, Thermo Fisher, CA) to mark cell membranes, streptavidin Alexa 647 (Thermo Fisher, CA) to mark sites of biotin accumulation (e.g. fatty acid biosynthesis in fat body, 1:500), and then washed four more times with SSCT, twice for five minutes and twice for 30 minutes. Samples were mounted in ProLong Gold Antifade Mountant (Thermo Fisher, CA), and imaged using a Zeiss LSM 880 with Airyscan fast at 10x and 40x.

### Tissue imaging and confocal microscopy

Beetles were dissected in PBS with a small amount of PBST added. Dissections were carried out in a similar fashion to those described under HCR. Samples were stored in ice cold PBST until all samples had been dissected. Samples were then fixed in 4% PFA with PBST for 25 minutes at room temperature. Samples were then quickly washed by exchanging the PBST three times, and then incubated with an anti-Engrailed 4D9 (1:5, DSHB, IA) mouse primary antibody in PBST overnight at 4°C. The following day, samples were rinsed quickly three times by replacing the PBST. Samples were then nutated 3 times for 20 minutes at room temperature with fresh PBST. Samples were then incubated with Alexa-488-Wheat Germ Agglutinin conjugate (WGA; 1:500), Hoechst 33342 (1:2000) to mark nuclei, and secondary antibody (Alexafluor-555 goat anti-mouse; 1:500, Thermo Fisher, CA) at room temperature for two hours. Samples were washed three times rapidly by replacing PBST, three 20 minute washes with PBST at room temp, and then were mounted in ProLong gold Antifade Mountant (Thermo Fisher, CA), and imaged using a Zeiss LSM 880 confocal microscope with Airyscan fast at 10x and 40x.

#### RNAi

Double-stranded RNA constructs were prepared as previously described^64^. Target sequences were cloned into a pCR2.1 TOPO vector (Thermo Fisher, CA) with primers containing T7 linkers. The following primers were used:

#### Dcor *oCYP4G*-1

F: 5’- TAATACGACTCACTATAGGGCACTCCCTGTCGGAACCTTGGA-3’

R: 5’-TAATACGACTCACTATAGGGTTGCGACATCCTCCACAGACGT-3’

#### Dcor *oCYP4G*-2

F: 5’-TAATACGACTCACTATAGGGACGTCTGTGGAGGATGTCGCAA-3’

R: 5’-TAATACGACTCACTATAGGGATCCAAAATCCCCGGACCCGAT-3’

#### Pson *oCYP4G*

F: 5’-TAATACGACTCACTATAGGGTCTTAGGATGTACCCACCAGTG-3’

R: 5’-TAATACGACTCACTATAGGGTGTCGCAATGCACTCGGTAT-3’ ***EGFP***:

F: 5’- TAATACGACTCACTATAGGGTCTTCTTCAAGGACGACGGCAACTAC -3’

R: 5’- TAATACGACTCACTATAGGGTTACTTGTACAGCTCGTCCATGCCGA -3’

A MEGAscript T7 Transcription kit (Thermo Fisher, CA) was used to synthesize dsRNA, which was subsequently cleaned using a MEGAclear Transcription cleanup kit (Thermo Fisher, CA), and quantified via NanoDrop (Thermo Fisher, CA). Target dsRNA and control dsRNA, targeting green fluorescent protein (GFP), were diluted in DEPC-treated PBS and green food dye to a concentration of ∼2mg/mL. Constructs were microinjected into third instar *Dalotia* larvae and adult *Platyusa*. *Dalotia* larvae were reared in 5cm Petri dishes with clean filter paper until they pupated and eclosed, after which they were fed frozen fruit flies until they were used in experiments. Injected *Dalotia* were used in experiments within 5-10 days of eclosing. *Platyusa* were maintained in Rubbermaid containers with a bed of packed damp coconut husk fiber and a diet of fruit flies. Beetles were sacrificed at 10 days and extracted in 100 microliters of hexane with a 10ng/microliter octadecane internal standard and analyzed via the CHC-GCMS method previously described. Total CHCs were compared between *oCYP4G-*KD and *GFP Platyusa* using a Welch’s t-test in R.

### qPCR of CHC pathway enzymes

Teneral *Sceptobius* were collected and reared as previously described. Three or four beetles were sampled each from zero, one, and three days post-eclosion as well as fully integrated beetles. All beetles were frozen at -80°C prior to RNA extraction. RNA was extracted and purified with the RNeasy purification kit (Qiagen, Germany). cDNA was then prepared using a Superscript III kit (Thermo Fisher, CA). The resulting cDNA was analyzed via qPCR. Briefly, 1 µL of cDNA template was combined with 6.5 µL of Luna® Universal qPCR master mix (New England BioLabs), 2 µL of combined forward and reverse primer (10µM, Integrated DNA Technologies, IA), and 10.5 µL of dH_2_O per well. Samples were run on an Applied Biosystems StepOnePlus Real Time PCR System (Thermo Fisher, CA) using the following conditions: 95°C for 1 minute, 40 cycles at 95° for 15 s, 59°C for 30s. Melt curve analysis was performed by holding for 15s at 95°C followed by 30s at 60°C and 15s at 95°C. Primers were designed for three *Sceptobius* CHC enzymes and two *Sceptobius* housekeeping genes, Ribosomal protein S3 and Ribosomal Protein L19.

The following primers were used:

#### Sceptobius ELO4

F: 5’-TTCGGTGCTGCAAGGATAA-3’

R: 5’-CCCAAGAGTACCACACAAGAG-3’

#### Sceptobius CYP4G1

F: 5’-ACCCAGCTGATATTGAGGTTATC-3’

R: 5’-CTGACCAAGAGACCATTTCCA-3’

#### Sceptobius Desat1

F: 5’-CGCTACAGCACTAACTTCACT-3’

R: 5’-ACACGACGCTTGACCATATC-3’

#### Sceptobius RPL19

F: 5’-GAATGTACCGCTACTGGTTTCT-3’

R: 5’-CAGCCTCTGTTATGCGATGT-3’

#### Sceptobius RPS3

F: 5’-GGTGTCGACGTAGTCGTTAATC-3’

R: 5’-CGAAGTAGTTGTGTCCGGTAAG-3’

Each sample was run in duplicate, and the geometric means of the technical replicates were determined. The ΔΔct calculation was then used to determine fold change of samples relative to the integrated condition. The difference between the cycle threshold for the genes of interest and the geometric mean of the two housekeeping genes was first calculated for each sample. From this difference in cycle threshold for each gene, the geometric mean of the cycle threshold difference in the integrated condition was subtracted. Multiplying the resulting values by negative one yielded the approximate fold change in expression for each time point compared to the integrated condition.

### Fluorescence quantification of CHC pathway enzyme expression levels

Teneral *Sceptobius* were collected and reared as previously described. Three beetles were sampled each from zero, one, two, three, and eight days post-eclosion. Four fully integrated *Sceptobius* were also collected. Due to the difficulty of obtaining larvae for the experiment, beetles were accumulated over the span of multiple weeks. As they were collected, samples were processed immediately via the previously mentioned HCR protocol up through the methanol dehydration step, at which point they were stored at -20°C. Once all samples had been collected, the remaining steps in the HCR process were carried out in parallel for all samples, with all beetles from the same time point being processed in the same tube. All sample tubes were incubated with identical probe and hairpin solution concentrations, targeting *Desat1*, *ELO4* and *CYP4G1*. Samples were mounted together and imaged on a Zeiss 880 confocal microscope (Zeiss, Germany) in Airyscan fast mode using identical imaging parameters. The integrated fluorescence signal was then calculated for all oenocytes in the resulting micrographs. Image stacks were converted to Tiffs and masks for oenocytes were determined using the *CYP4G1* channel and the python package Cellpose v.2.0.5^154^ in Python v3.8.13. Image stacks were first projected along the z-axis by summing over all slices in the stack and then cells were identified with the ‘cyto’ model in Cellpose using a diameter of 35, flow threshold of 0.1, minimum size of 200, cell probability threshold of 0.8, and interp and neg_avg set to true. The resulting oenocyte cell masks were used to filter the *CYP4G1* channel by calculating the percentage of the z-projected frame occupied by all cell masks, and then setting a pixel intensity threshold that excludes the same percentage of voxels across the *CYP4G1* channel slices. The threshold mask determined with the *CYP4G1* channel was then applied to the *ELO4* and *Desat1* channels. Next, the cell masks were applied to each filtered channel, integrating over the fluorescence intensity in each slice to recover the total fluorescence intensity in each oenocyte, for the three channels.

### Aggression analysis

Wild caught *Sceptobius*, *Liometopum*, and *Platyusa* were transported to the lab as previously described and maintained until they were used in behavior experiments. *Liometopum* and *Sceptobius* were used in behavioral arenas within a week of collecting to avoid shifts in CHC profiles that arise from the standard lab feeding setup. *Sceptobius* were coated with *Platyusa* CHCs by placing *Sceptobius* in 5 mL vials with 7-8 *Platyusa* for 2 hours. The number of *Platyusa* was sufficient to fill the entire bottom of the vial, ensuring that *Sceptobius* was in contact with *Platyusa* for the entire 2 hour period, regardless of grooming status. Control *Sceptobius* were placed in empty vials for 2 hours. Following the coating procedure, *Sceptobius* was loaded into a behavioral arena with three *Liometopum* collected contemporaneously from the same colony as the beetle. The behavioral arena was constructed from acrylic, with a 5cm diameter and a sliding door to allow acclimation of the beetle and ants in the arena prior to being introduced to each other. Beetles and ants were loaded into the arena unanesthetized, to avoid changes in behavior that arise from CO_2_ or cold anesthesia^155–157^. After a 2 minute acclimation period, the sliding door was opened, allowing the beetle and the ants to interact for a 5 minute interaction period. Behavior was recorded on a FLIR camera (BFS-U3-16S2C-CS: 1.6 MP) with an InfiniGage lens (Infinity Photo-Optical, CO), at 60hz.

After each run, *Sceptobius* and the three *Liometopum* CHCs were analyzed via GCMS. Each beetle was extracted in 10 microliters of hexane for 20 minutes and three microliters of extract were analyzed. *Liometopum* samples were extracted in 70 microliters of hexane for 20 minutes and two microliters were analyzed. CHCs were run on the GCMS, identified, and integrated as described previously. Scoring of behavioral interactions was carried out in BORIS v. 8.20.1^158^. A simple ethogram consisting of biting, mandible flaring, and neutral interaction (any instance where no biting or mandible flaring was observed) was used. Interactions were scored any time that any ant’s antenna made contact with *Sceptobius*. An aggression index was calculated as the sum of mandible flaring events and two times the number of biting events, divided by the sum of mandible flaring events, two times the number of biting events, and neutral events. Aggression values can thus take any value between zero, i.e. none of the interactions were aggressive, and one, i.e. every interaction was aggressive.

### Behavioral analysis of beetle-ant physical interactions

A multi-well behavior arena was constructed from 1/8^th^ inch infrared transparent acrylic (Plexiglass IR acrylic 3143) and was backlit with infrared LED panel. To ensure that runs were conducted in complete darkness, the arena was placed inside of an incubator in a dark room. The arena was 4 cm in diameter, split into multiple chambers to give the beetles places to hide from their paired ant. Small, ∼5 mm diameter, pieces of damp filter paper were added to each well to increase humidity within wells. Single *Sceptobius lativentris, Dalotia coriaria,* and *Platyusa sonomae* were cold anesthetized, and then loaded into individual arena wells, along with one cold anesthetized *Liometopum occidentale*. Beetles were recorded at 1 hz for 24 hours using a FLIR camera (BFS-U3-51S5M-C: 5.0 MP) with a Pentax 12mm 1:1.2 TV lens (Ricoh, FL-HC1212B-VG). Videos were cropped down to individual wells and analyzed with Deeplabcut v2.0.6.2^159,160^. Distinct network models, using a default ResNet50 network, were trained for each beetle, consisting of two labels for the head and abdominal tip of both *Liometopum* and *Platyusa*, and one label for the head of *Sceptobius* and *Dalotia*. The networks were trained on ∼500 labeled frames for the *Platyusa* model, ∼150 frames for the *Sceptobius* model, and ∼50 frames for the *Dalotia* model. All of the models were trained over multiple iterations and the *Dalotia* model used the training weights from the final *Sceptobius* model as its initial weights. The training and test errors were 1.27 and 0.92 pixels for the *Platyusa* model, 1.03 and 2.79 pixels for the *Sceptobius* model, and 2.13 and 3.55 pixels for the *Dalotia* model. The three networks were used to analyze all of the videos of their respective beetles.

Position data was filtered for frames in which the ant prediction is likely accurate by calculating the length of each ant, and then choosing an upper threshold that excludes outliers for all ants collected in a given behavioral run. Then an ellipse was drawn around the ant for each labelled video frame, such that the sum of the distances between any point on the perimeter of the ellipse and the head and abdominal tip of the ant summed to 1.4 times the ant length in that frame. If the beetle’s head, or head or tail for *Platyusa*, fell within the ellipse for that frame, the beetle was considered to be touching the ant. The percent of frames in which the beetle was touching the ant was then calculated for each well. Differences in the touching percent between the three beetles were compared using an ANOVA with a Tukey *post hoc* test in R.

#### Desiccation Analysis

Desiccation resistance was measured in *Sceptobius* using an acrylic desiccation arena. The arena consisted of two sheets of 3/8 inch acrylic both containing six wells with a diameter of one centimeter each. The two sets of wells sandwiched a piece of filter paper, which acted as the floor for the beetles, located in the upper set of wells. The lower set of wells was used to introduce wetted circles of filter paper as a humidity source. The well setup was then sandwiched between two sets of IR passthrough acrylic, creating a dark behavior arena. The desiccation arena was lit from below by LED lights, which emitted partially in the IR range.

To test the effect of superphysiological levels of CHCs on *Sceptobius,* beetles were coated in *Liometopum* CHCs, which were extracted and fractionated previously^89^. Briefly, surface chemicals were extracted from tens of thousands of *Liometopum*, and were subsequently separated via vacuum flash chromatography and eluted with hexane and cyclohexane. The resulting purified CHC extracts contained approximately 250 ant equivalents per milliliter. Roughly 25 ant equivalents of *Liometopum* CHCs were then applied to the inside of a 5mL glass vial and the hexane was evaporated off under a stream of N_2_. This process resulted in a thin coating of *Liometopum* CHCs on the inner surface of the vial. *Sceptobius* were collected at multiple nest-sites, and were pooled for each run. Beetles were either placed in the CHC coated vial, or in a blank control vial, for an hour. *Sceptobius* in the CHC coated vial moved around enough to transfer CHCs from the walls of the vial to their cuticle Groups of five beetles were then loaded into each well in the arena. Beetles were run in groups to avoid confounding effects from social isolation. Desiccation trials were run at room temperature under a FLIR camera (BFS-U3-04S2M-CS: 0.4 MP) with an InfiniGage lens (Infinity Photo-Optical, CO) running at 1 hz. Recordings were collected until all beetles in the desiccation arena were dead, typically 20-40 hours.

For runs with higher humidity, 10 pieces of hole punched filter paper were placed in the lower chamber below each well and 50µl of water were added to the pieces of filter paper. To replicate the effects of CHC addition on another beetle species, we knocked down *CYP4G1* in *Dalotia coriaria,* using the RNAi protocol described above. Control beetles were injected with dsRNA for *GFP*, and thus had a species typical CHC profile. We then took the *CYP4G* KD *Dalotia* and coated them in ant CHCs, as described above. *GFP* RNAi *Dalotia*, *CYP4G* RNAi *Dalotia*, and *CYP4G* RNAi *Dalotia* with *Liometopum* CHCs were then run in the desiccation arena. After each run, *Dalotia* were extracted in 70 microliters of hexane containing an internal octadecane standard and analyzed with the CHC-GCMS method describe above to verify that the knockdowns were successful. Beetles for which the wild type CHC profile was detectible were excluded from downstream analysis.

### Sceptobius and Platyusa isolation from Liometopum

*Sceptobius* were maintained in a container of damp coconut fiber in an incubator at 24°C and 90-95% relative humidity. Control beetles were kept in similar containers containing 12 *Liometopum* workers. Boxes were monitored twice daily for beetle death. CHC amounts were quantified for living or recently deceased *Sceptobius* at 48 and 72 hours of isolation by extracting beetles in 10 microliters of hexane containing 1ng/microliter of C18 for 20 minutes followed by GCMS analysis using the previously described CHC-GCMS method. Wild caught *Platyusa* were reared for a month on a diet of frozen *Drosophila melanogaster* in the absence of *Liometopum occidentale*. *Platyusa* and *Drosophila* were then extracted in 70 microliters of hexane containing 10 ng/microliter C18 for 20 minutes and run on the GCMS using the previously described CHC GCMS method. CHC profiles of the *Drosophila* fed *Platyusa* were compared to CHC profiles of *Platyusa* fed frozen *Liometopum occidentale* for the same period.

---

**Table S1.** Average L occidentale, P sonomae, S lativentris, and L newtonarum CHCs.

|  | **RI** | ***Liometopum*** | ***Sceptrubius*** | ***Platyusa*** | ***Liometoxenus*** |
| --- | --- | --- | --- | --- | --- |
| C23 | 2300 | 0.42±0.48 | 0.2±0.13 | 0.18±0.13 | 0.45±0.14 |
| 11;9meC23 | 2332 | 0.04±0.08 | 0.003±0.01 | 0.01±0.01 | 0.04±0.05 |
| 3meC23 | 2372 | 0.02±0.04 | 0.003±0.02 | – | 0.03±0.04 |
| C24 | 2400 | 0.13±0.11 | 0.16±0.11 | 0.25±0.08 | 0.13±0.05 |
| xmeC24 | 2433 | 0.03±0.05 | 0.02±0.05 | 0.02±0.02 | 0.06±0.04 |
| C25:1 (2) | 2477 | 0.1±0.13 | 0.01±0.07 | 0.01±0.02 | 0.16±0.08 |
| C25 | 2500 | 6.05±3.11 | 3.59±2.04 | 17.77±6.03 | 5.17±2 |
| 13;11;9meC25 | 2532 | 1.9±2.25 | 0.46±0.19 | 0.66±0.68 | 1.79±0.76 |
| 7meC25 | 2539 | 0.05±0.06 | 0.05±0.04 | 0.03±0.03 | 0.07±0.03 |
| 5meC25 | 2548 | 0.36±0.29 | 0.14±0.08 | 0.26±0.21 | 0.67±0.24 |
| dimeC25 (1) | 2561 | 0.18±0.28 | 0.06±0.09 | 0.17±0.14 | 0.18±0.08 |
| 3meC25 | 2573 | 0.98±0.79 | 0.35±0.55 | 6.84±2.78 | 1.02±0.26 |
| dimeC25 (2) | 2580 | 0.34±0.37 | 0.14±0.2 | 0.11±0.1 | 0.43±0.12 |
| C26 | 2600 | 0.33±0.12 | 0.28±0.13 | 0.35±0.13 | 0.18±0.09 |
| dimeC25 (3) | 2605 | 0.31±0.41 | 0.08±0.06 | 0.11±0.1 | 0.36±0.12 |
| 13;12;11;10meC26 | 2631 | 0.22±0.25 | 0.09±0.07 | 0.07±0.07 | 0.21±0.08 |
| C27:2 (Lnew) | 2653 | – | – | – | 0.53±0.74 |
| 4meC26 | 2657 | – | – | 0.08±0.02 | – |
| C27:2 | 2677 | – | – | 1.41±1.11 | – |
| C27:1 (3) | 2678 | 1.42±0.96 | 0.7±0.22 | 0.94±0.75 | 2.29±0.53 |
| C27:1 (4) | 2685 | – | – | 1.24±0.94 | – |
| C27 | 2700 | 4.82±1.67 | 3.15±1.25 | 1.12±0.39 | 1.42±0.6 |
| 13;11;9meC27 | 2730 | 3.31±2.46 | 1.96±0.61 | 0.96±0.73 | 3.25±1.08 |
| 7meC27 | 2739 | 0.29±0.18 | 0.19±0.05 | 0.06±0.05 | 0.23±0.09 |
| 5meC27 | 2748 | 0.41±0.16 | 0.43±0.13 | 0.36±0.12 | 1.28±0.69 |
| dimeC27 (1) | 2757 | 0.23±0.34 | 0.09±0.07 | 0.18±0.11 | 0.2±0.09 |
| dimeC27 (2) | 2764 | 0.21±0.29 | 0.1±0.07 | 0.21±0.57 | 0.25±0.1 |
| dimeC27 (3) | 2768 | 0.45±0.61 | 0.15±0.09 | 0.05±0.13 | 0.4±0.17 |
| 3meC27 | 2772 | 1.11±0.43 | 0.82±0.23 | 0.33±0.08 | 0.82±0.2 |
| dimeC27 (4) | 2777 | 0.34±0.5 | – | 0.31±0.17 | – |
| C28:1 | 2778 | 0.5±0.38 | 0.65±0.18 | 0.01±0.06 | 1.04±0.16 |
| dimeC27 (p1) | 2785 | – | – | 0.08±0.03 | – |
| dimeC27 (p2) | 2790 | – | – | 0.09±0.05 | – |
| C28 | 2800 | 0.51±0.25 | 0.44±0.2 | 0.06±0.03 | 0.1±0.03 |
| dimeC27 (5) | 2803 | 0.47±0.46 | 0.16±0.11 | 0.16±0.11 | 0.75±0.32 |
| 14;13;12;10meC28 | 2829 | 0.69±0.38 | 0.11±0.27 | 0.16±0.08 | 0.75±0.46 |
| C29:2 (1) | 2861 | 0.23±0.18 | 0.19±0.12 | 9.46±4.18 | 0.42±0.21 |
| C29:2 (2) | 2869 | 0.39±0.24 | 0.69±0.35 | 0.16±0.09 | 3.32± |
| C29:1 (1) | 2871 | 0.54±0.27 | 1.46±0.56 | 0.26±0.07 | 0.51±0.07 |
| C29:1n9 | 2879 | 22.96±10.33 | 30.18±4.68 | 10.21±2.98 | 30.61±5.42 |
| C29:1n7 | 2886 | 0.18±0.09 | 0.22±0.09 | 4.1±1.16 | 0.3±0.09 |
| C29 | 2900 | 8.5±4.4 | 6.94±2.15 | 0.77±0.39 | 1.13±0.46 |
| 15;13;11;9meC29 | 2930 | 4.83±2.3 | 5.02±1.08 | 1.91±0.55 | 3.96±1 |
| 7meC29 | 2940 | 1.1±0.31 | 1.33±0.76 | 0.21±0.07 | 0.95±0.3 |
| 5meC29 | 2949 | 0.65±0.15 | 0.64±0.14 | 0.68±0.26 | 0.99±0.5 |
| dimeC29 (1) | 2954 | 0.34±0.26 | 0.18±0.29 | 3.02±0.79 | 0.26±0.1 |
| dimeC29 (2) | 2961 | 0.44±0.28 | 0.34±0.16 | 0.3±0.11 | 0.48±0.15 |
| dimeC29 (3) | 2968 | 0.8±0.59 | 0.56±0.18 | 0.06±0.16 | 0.72±0.27 |
| 3meC29 | 2975 | 1.85±0.69 | 1.88±0.44 | 0.45±0.37 | 0.55±0.16 |
| C30:1 | 2980 | 0.65±0.56 | 1±0.29 | – | 0.69±0.12 |
| dimeC29 (4) | 2981 | 0.16±0.24 | – | 1.34±0.75 | – |
| dimeC29 (p1) | 2988 | – | – | 0.7±0.28 | – |
| dimeC29 (5) | 3001 | 0.79±0.22 | 0.92±0.29 | 0.22±0.05 | 0.41±0.11 |
| 15;14;13;12;10meC30 | 3029 | 0.56±0.22 | 0.6±0.16 | 0.31±0.08 | 0.34±0.12 |
| C31:3 | 3037 | – | – | – | 3.53±4.53 |
| C31:2 (1) | 3056 | 0.81±0.45 | 0.84±0.23 | 10.88±3.49 | 1.57±0.52 |
| C31:2 (2) | 3060 | 3.7±2.29 | 4.04±1.34 | 0.94±0.41 | 7.33±1.98 |
| C31:2 (3) | 3066 | 0.13±0.09 | 0.15±0.09 | 0.1±0.04 | 0.22±0.07 |
| C31:1 (1) | 3075 | 0.06±0.12 | – | – | 0.25±0.08 |
| C31:2 (4) | 3077 | 0.27±0.24 | 0.34±0.6 | 0.38±0.1 | – |
| C31:1 (3) | 3082 | 1.9±1.11 | 2.27±0.6 | 0.29±0.2 | 1.61±0.44 |
| C31:1 (4) | 3090 | – | – | 0.14±0.08 | – |
| C31 | 3100 | 1.22±1.1 | 1.03±0.39 | 0.07±0.04 | 0.14±0.06 |
| 15;13;11;9meC31 | 3131 | 4.62±1.41 | 5.6±0.73 | 1.29±0.4 | 3.05±0.57 |
| 5meC31 | 3150 | 0.34±0.12 | 0.23±0.08 | 0.16±0.1 | 0.16±0.04 |
| dimeC31 (1) | 3153 | 0.5±0.32 | 0.54±0.16 | 2.54±0.89 | 0.45±0.11 |
| dimeC31 (2) | 3164 | 1.18±0.64 | 1.38±0.49 | 0.52±0.3 | 0.77±0.22 |
| dimeC31 (3) | 3167 | 0.21±0.16 | 0.24±0.13 | 0.23±0.36 | 0.17±0.12 |
| 3meC31 | 3175 | 0.57±0.36 | 0.49±0.08 | 0.73±0.58 | 0.19±0.07 |
| dimeC31 (p1) | 3187 | – | – | 0.45±0.84 | – |
| C33:3 | 3231 | – | – | – | 1.56±1.76 |
| 33:2 (1) | 3252 | 0.01±0.06 | – | 2.73±3.02 | – |
| 33:2 (2) | 3260 | 0.04±0.18 | – | 0.64±0.75 | – |
| 17;15;13;11;9meC33 | 3328 | 4.37±1.43 | 5.79±1.25 | 0.77±0.3 | 2.83±0.66 |
| dimeC33 (1) | 3357 | 3.53±1.67 | 3.57±0.9 | 2.19±0.68 | 2.44±0.74 |
| dimeC33 (2) | 3374 | 0.48±0.15 | 0.6±0.17 | 0.42±0.18 | 0.38±0.14 |
| C35:2 (1) | 3452 | – | – | 0.71±0.63 | – |
| C35:2 (2) | 3458 | – | – | 0.14±0.14 | – |
| 17;15;13;11meC35 | 3525 | 2.44±0.9 | 3.38±0.76 | 1.07±0.38 | 1.54±0.35 |
| dimeC35 | 3553 | 2.42±1.24 | 2.77±0.88 | 3.81±1.73 | 1.93±0.55 |

**Table S2.** CHC Biosynthesis Enzymes in *Platyusa*.

| **Gene** | **Pson Orthologs** |
| --- | --- |
| ELO4 | Pson_TRINITY_DN782_c0_g1_i3.p1*, Pson_TRINITY_DN782_c0_g1_i4.p1 |
| ELO1 | none |
| ELO2 | Pson_TRINITY_DN9471_c0_g1_i1.p2 |
| ELO3 | Pson_TRINITY_DN2473_c1_g1_i1.p1 |
| ELO5 | Pson_TRINITY_DN2473_c0_g1_i1.p1 |
| CyP4G1 | Pson_TRINITY_DN2722_c0_g2_i1.p1* |
| desat1 | Pson_TRINITY_DN257_c2_g1_i1.p1* |
| FAS1 | Pson_TRINITY_DN3393_c0_g1_i4.p1, Pson_k119_722322_gene2a |
| FAS2 | Pson_TRINITY_DN33_c0_g1_i7.p1, Pson_TRINITY_DN3888_c0_g1_i1.p1 |
| FAS3 | Pson_TRINITY_DN465_c0_g1_i4.p1* |
| FAR1 | Pson_TRINITY_DN7112_c0_g1_i9.p1,<br>Pson_TRINITY_DN8317_c1_g1_i19.p1 |
| FAR2 | Pson_TRINITY_DN1026_c0_g1_i10.p1 |
| FAR3 | Pson_TRINITY_DN5094_c0_g1_i4.p1 |
| FAR4 | Pson_TRINITY_DN4620_c0_g1_i6.p1*,<br>Pson_TRINITY_DN4620_c0_g2_i13.p1,<br>Pson_k119_936437_483757_1086134_248186_1114230RC_gene4a |
| desat2 | Pson_TRINITY_DN1356_c1_g1_i8.p1 |
| desat4 | Pson_TRINITY_DN4819_c0_g1_i1.p1 |
| desat3 | Pson_TRINITY_DN440_c0_g1_i11.p2 |
| desat5 | Pson_TRINITY_DN7573_c0_g1_i6.p1, Pson_TRINITY_DN7573_c0_g1_i8.p1 |
| ELO6 | Pson_TRINITY_DN10704_c0_g2_i4.p1 |
| ELO7 | Pson_TRINITY_DN1185_c0_g1_i19.p1 |
| ELO8 | Pson_TRINITY_DN3893_c0_g1_i1.p1 |
| ELO9 | Pson_TRINITY_DN6750_c0_g1_i1.p1 |
| ELO10 | Pson_TRINITY_DN6520_c0_g1_i8.p1 |
| FAR5 | Pson_TRINITY_DN155_c97_g1_i7.p1 |
| FAR6 | Pson_TRINITY_DN32631_c0_g1_i1.p1 |
| FAR7 | Pson_k119_245651_360072_58654_403476_814044RC_958313RC_gene1a |
| ELO11 | Pson_TRINITY_DN112378_c0_g1_i1.p1 |
| CPR | Pson_TRINITY_DN753_c1_g2_i4.p1 |
| FAR8 | Pson_TRINITY_DN2744_c0_g1_i5.p1,<br>Pson_TRINITY_DN2254_c0_g1_i22.p1 |
| ELO12 | Pson_TRINITY_DN5856_c0_g1_i1.p1 |

\*denotes sequences used for HCR probe development

**Table S3.**
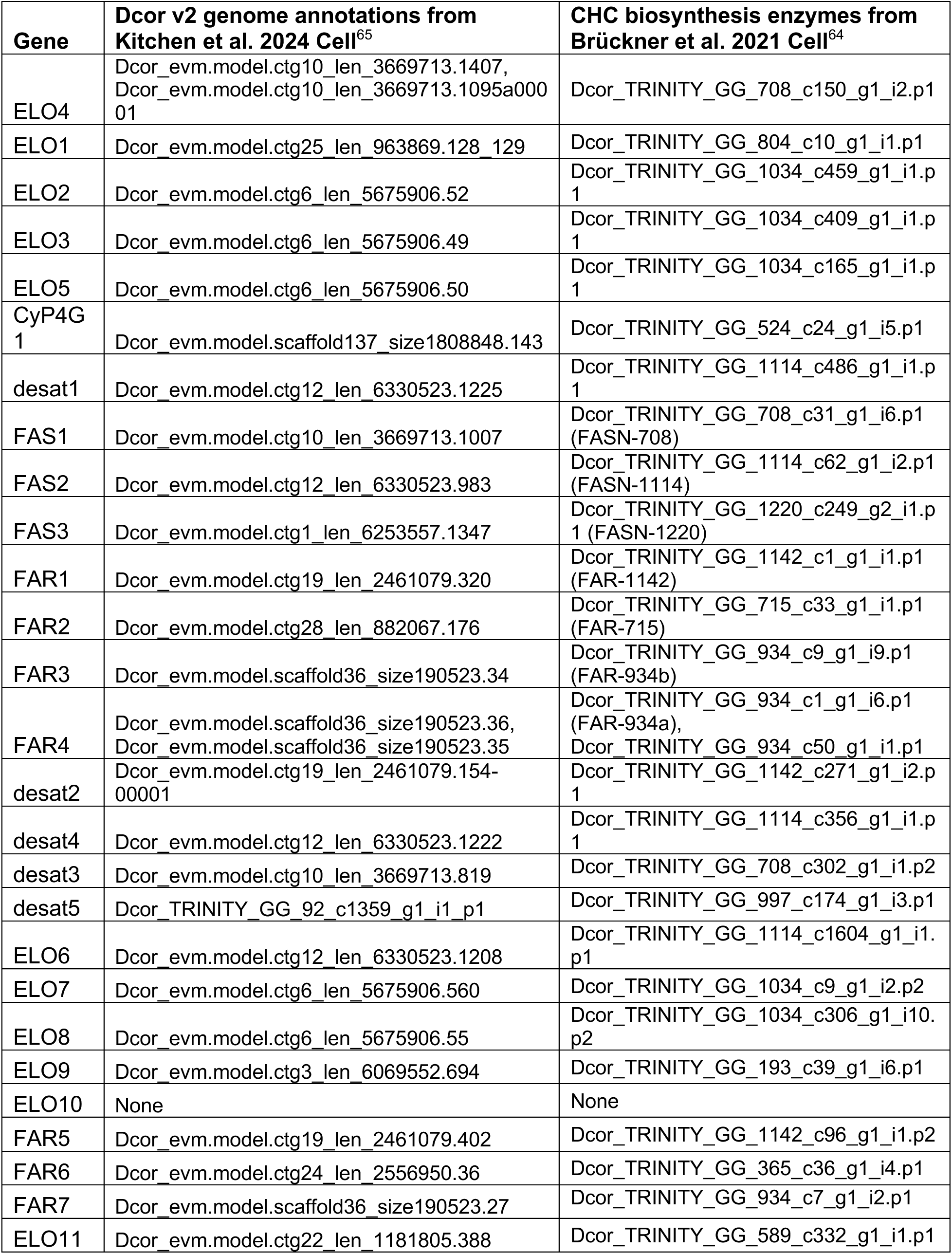
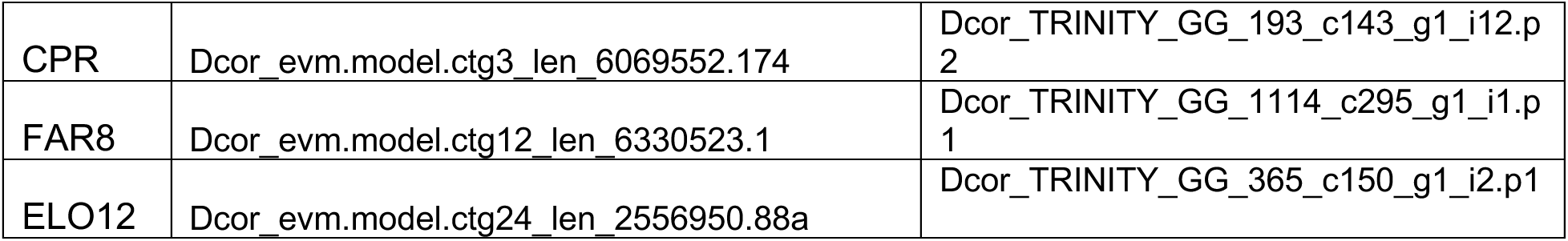
CHC Biosynthesis Enzyme Orthologs in *Dalotia*.

| **Gene** | **Dcor v2 genome annotations from Kitchen et al. 2024 Cell<sup>65</sup>** | **CHC biosynthesis enzymes from Brückner et al. 2021 Cell<sup>64</sup>** |
| --- | --- | --- |
| ELO4 | Dcor_evm.model.ctg10_len_3669713.1407,<br>Dcor_evm.model.ctg10_len_3669713.1095a00001 | Dcor_TRINITY_GG_708_c150_g1_i2.p1 |
| ELO1 | Dcor_evm.model.ctg25_len_963869.128_129 | Dcor_TRINITY_GG_804_c10_g1_i1.p1 |
| ELO2 | Dcor_evm.model.ctg6_len_5675906.52 | Dcor_TRINITY_GG_1034_c459_g1_i1.p1 |
| ELO3 | Dcor_evm.model.ctg6_len_5675906.49 | Dcor_TRINITY_GG_1034_c409_g1_i1.p1 |
| ELO5 | Dcor_evm.model.ctg6_len_5675906.50 | Dcor_TRINITY_GG_1034_c165_g1_i1.p1 |
| Cyp4G1 | Dcor_evm.model.scaffold137_size1808848.143 | Dcor_TRINITY_GG_524_c24_g1_i5.p1 |
| desat1 | Dcor_evm.model.ctg12_len_6330523.1225 | Dcor_TRINITY_GG_1114_c486_g1_i1.p1 |
| FAS1 | Dcor_evm.model.ctg10_len_3669713.1007 | Dcor_TRINITY_GG_708_c31_g1_i6.p1 (FASN-708) |
| FAS2 | Dcor_evm.model.ctg12_len_6330523.983 | Dcor_TRINITY_GG_1114_c62_g1_i2.p1 (FASN-1114) |
| FAS3 | Dcor_evm.model.ctg1_len_6253557.1347 | Dcor_TRINITY_GG_1220_c249_g2_i1.p1 (FASN-1220) |
| FAR1 | Dcor_evm.model.ctg19_len_2461079.320 | Dcor_TRINITY_GG_1142_c1_g1_i1.p1 (FAR-1142) |
| FAR2 | Dcor_evm.model.ctg28_len_882067.176 | Dcor_TRINITY_GG_715_c33_g1_i1.p1 (FAR-715) |
| FAR3 | Dcor_evm.model.scaffold36_size190523.34 | Dcor_TRINITY_GG_934_c9_g1_i9.p1 (FAR-934b) |
| FAR4 | Dcor_evm.model.scaffold36_size190523.36,<br>Dcor_evm.model.scaffold36_size190523.35 | Dcor_TRINITY_GG_934_c1_g1_i6.p1 (FAR-934a),<br>Dcor_TRINITY_GG_934_c50_g1_i1.p1 |
| desat2 | Dcor_evm.model.ctg19_len_2461079.154-00001 | Dcor_TRINITY_GG_1142_c271_g1_i2.p1 |
| desat4 | Dcor_evm.model.ctg12_len_6330523.1222 | Dcor_TRINITY_GG_1114_c356_g1_i1.p1 |
| desat3 | Dcor_evm.model.ctg10_len_3669713.819 | Dcor_TRINITY_GG_708_c302_g1_i1.p2 |
| desat5 | Dcor_TRINITY_GG_92_c1359_g1_i1.p1 | Dcor_TRINITY_GG_997_c174_g1_i3.p1 |
| ELO6 | Dcor_evm.model.ctg12_len_6330523.1208 | Dcor_TRINITY_GG_1114_c1604_g1_i1.p1 |
| ELO7 | Dcor_evm.model.ctg6_len_5675906.560 | Dcor_TRINITY_GG_1034_c9_g1_i2.p2 |
| ELO8 | Dcor_evm.model.ctg6_len_5675906.55 | Dcor_TRINITY_GG_1034_c306_g1_i10.p2 |
| ELO9 | Dcor_evm.model.ctg3_len_6069552.694 | Dcor_TRINITY_GG_193_c39_g1_i6.p1 |
| ELO10 | None | None |
| FAR5 | Dcor_evm.model.ctg19_len_2461079.402 | Dcor_TRINITY_GG_1142_c96_g1_i1.p2 |
| FAR6 | Dcor_evm.model.ctg24_len_2556950.36 | Dcor_TRINITY_GG_365_c36_g1_i4.p1 |
| FAR7 | Dcor_evm.model.scaffold36_size190523.27 | Dcor_TRINITY_GG_934_c7_g1_i2.p1 |
| ELO11 | Dcor_evm.model.ctg22_len_1181805.388 | Dcor_TRINITY_GG_589_c332_g1_i1.p1 |
| CPR | Dcor_evm.model.ctg3_len_6069552.174 | Dcor_TRINITY_GG_193_c143_g1_i12.p<br>2 |
| FAR8 | Dcor_evm.model.ctg12_len_6330523.1 | Dcor_TRINITY_GG_1114_c295_g1_i1.p<br>1 |
| ELO12 | Dcor_evm.model.ctg24_len_2556950.88a | Dcor_TRINITY_GG_365_c150_g1_i2.p1 |

**Table S4.** CHC Biosynthesis Enzymes in *Sceptobius*.

| **Gene** | **Slat Orthologs** |
| --- | --- |
| ELO4 | Slat_TRINITY_DN165_c1_g1_i28.p1* |
| ELO1 | Slat_TRINITY_DN20689_c0_g1_i7.p1 |
| ELO2 | Slat_evm.model.hic_scaffold_2_pilon_pilon.379-00001 |
| ELO3 | Slat_TRINITY_DN9690_c1_g1_i9.p1,<br>Slat_evm.model.hic_scaffold_2_pilon_pilon.373,<br>Slat_evm.model.hic_scaffold_2_pilon_pilon.374 |
| ELO5 | Slat_TRINITY_DN54_c0_g2_i2.p1 |
| CyP4G1 | Slat_TRINITY_DN299_c0_g1_i3.p1* |
| desat1 | Slat_TRINITY_DN897_c0_g1_i37.p1* |
| FAS1 | Slat_TRINITY_DN902_c7_g1_i101.p1 |
| FAS2 | None |
| FAS3 | Slat_TRINITY_DN18025_c1_g1_i4.p1,<br>Slat_TRINITY_DN508_c3_g1_i5.p1,<br>Slat_evm.model.hic_scaffold_3_pilon_pilon.1657*,<br>Slat_evm.model.hic_scaffold_3_pilon_pilon.1761,<br>Slat_evm.model.hic_scaffold_5_pilon_pilon.1559 |
| FAR1 | Slat_TRINITY_DN2352_c0_g1_i3.p1,<br>Slat_TRINITY_DN4862_c0_g1_i12.p1 |
| FAR2 | Slat_TRINITY_DN454_c4_g1_i11.p1,<br>Slat_TRINITY_DN1701_c0_g1_i42.p1 |
| FAR3 | Slat_TRINITY_DN4373_c0_g1_i4.p1 |
| FAR4 | Slat_evm.model.hic_scaffold_9_pilon_pilon.512* |
| desat2 | Slat_TRINITY_DN1161_c4_g1_i2.p1 |
| desat4 | Slat_TRINITY_DN557_c1_g2_i1.p1 |
| desat3 | Slat_evm.model.hic_scaffold_10_pilon_pilon.1681 |
| desat5 | None |
| ELO6 | Slat_TRINITY_DN15725_c0_g1_i12.p1 |
| ELO7 | Slat_TRINITY_DN15611_c0_g1_i2.p1 |
| ELO8 | Slat_TRINITY_DN3691_c0_g1_i1.p1 |
| ELO9 | Slat_TRINITY_DN1299_c0_g2_i2.p1 |
| ELO10 | Slat_TRINITY_DN28641_c0_g1_i3.p1 |
| FAR5 | Slat_TRINITY_DN35465_c0_g1_i4.p1,<br>Slat_TRINITY_DN710_c1_g1_i5.p1 |
| FAR6 | Slat_evm.model.hic_scaffold_9_pilon_pilon.915 |
| FAR7 | Slat_evm.model.hic_scaffold_9_pilon_pilon.697,<br>Slat_evm.model.hic_scaffold_9_pilon_pilon.1117,<br>Slat_evm.model.hic_scaffold_4_pilon_pilon.2085 |
| ELO11 | Slat_TRINITY_DN2208_c0_g1_i15.p1 |
| CPR | Slat_TRINITY_DN4772_c0_g1_i1.p1 |
| FAR8 | Slat_TRINITY_DN4813_c4_g1_i13.p1 |
| ELO12 | Slat_TRINITY_DN2842_c0_g1_i11.p3,<br>Slat_TRINITY_DN7230_c0_g1_i20.p1,<br>Slat_evm.model.hic_scaffold_9_pilon_pilon.77 |

\*denotes sequences used for HCR probe development

## Supplemental figure legends

**Figure S1.**
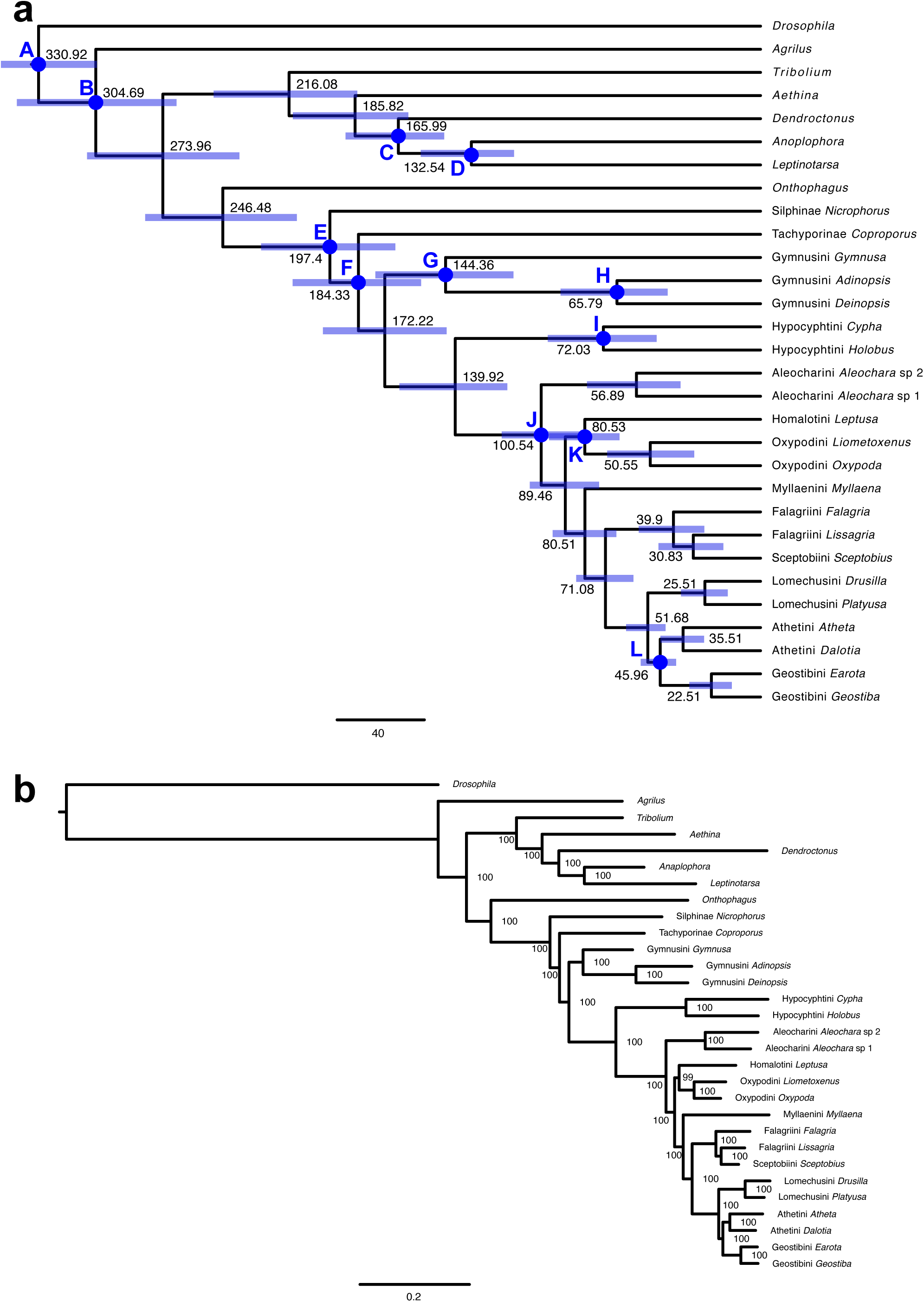
Additional species trees. **a:** A time calibrated species tree generated with the approximate Bayesian method in MCMCTree. Divergence times are listed in millions of years (Ma), blue bars represent 95% credibility intervals on divergence times, and labeled blue dots specify fossil calibration points, listed in methods. **b:** A maximum likelihood species tree, with bootstrap support listed at each node.

**Figure S2.**
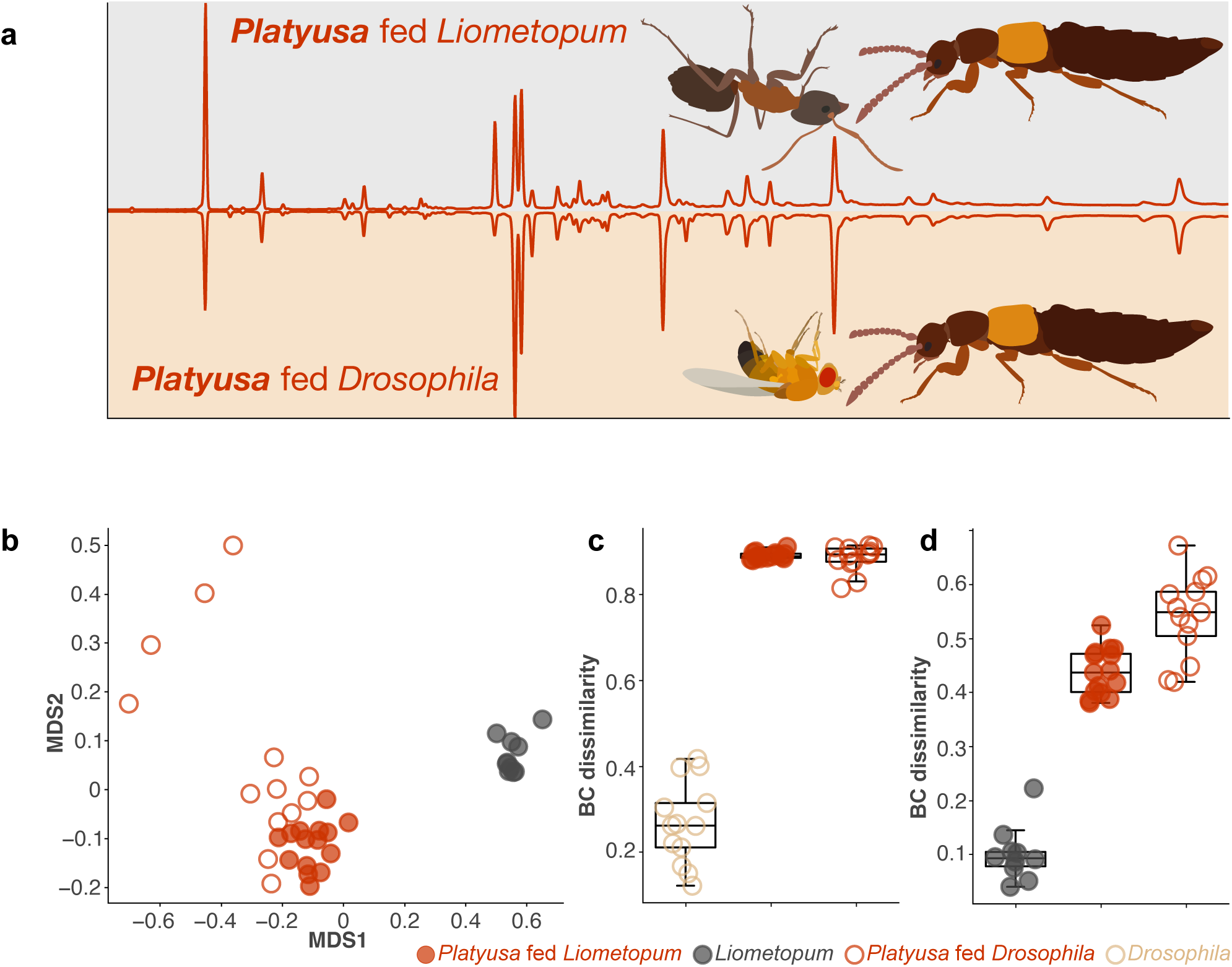
Platyusa CHCs after isolation. **a:** Representative GC traces showing the CHC profile of *Platyusa* fed *Liometopum* or *Drosophila* for one month. **b:** NMDS ordination plot of pairwise Bray Curtis dissimilarity measurements of *Liometopum*, *Platyusa* fed *Liometopum* for one month, and *Platyusa* fed *Drosophila* for one month (stress=0.0859) **c:** Bray Curtis dissimilarity between the two *Platyusa* treatments and the average *Drosophila* CHC profile. *Drosophila* samples were also compared to the average *Drosophila* profile. **d:** Bray Curtis dissimilarity between the two *Platyusa* treatments and the average *Liometopum* CHC profile. *Liometopum* samples were also compared to the average *Liometopum* profile.

**Figure S3.**
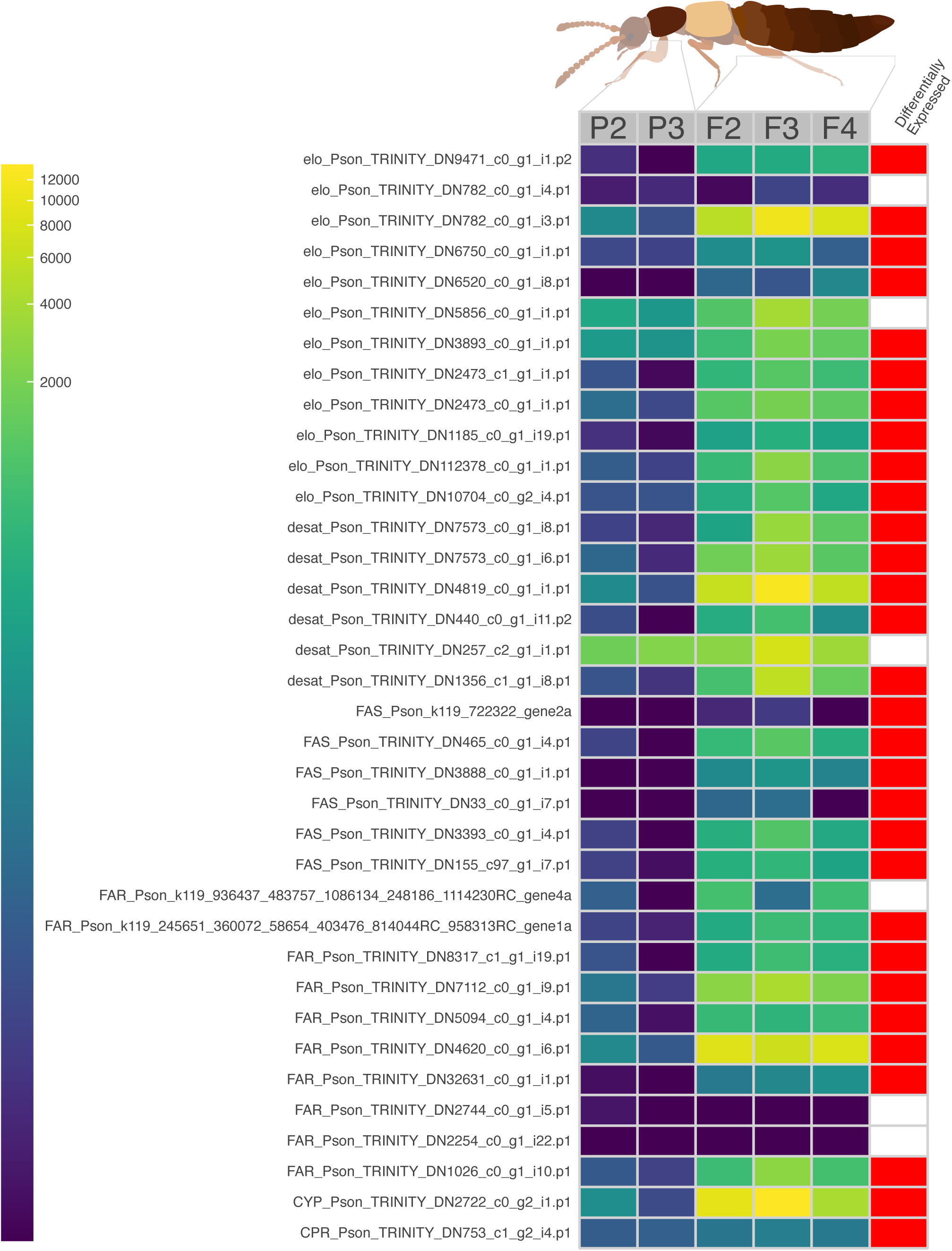
CHC biosynthesis enzyme expression in *Platyusa sonomae*. Heatmap of CHC enzyme expression, in transcripts per million (TPM) for control tissue (pronotum, P) and test tissue (fat body, F). Numbers next to the treatment group correspond to the id of the single beetle dissected for that sample. Differentially expressed transcripts are shown in red.

**Figure S4.**
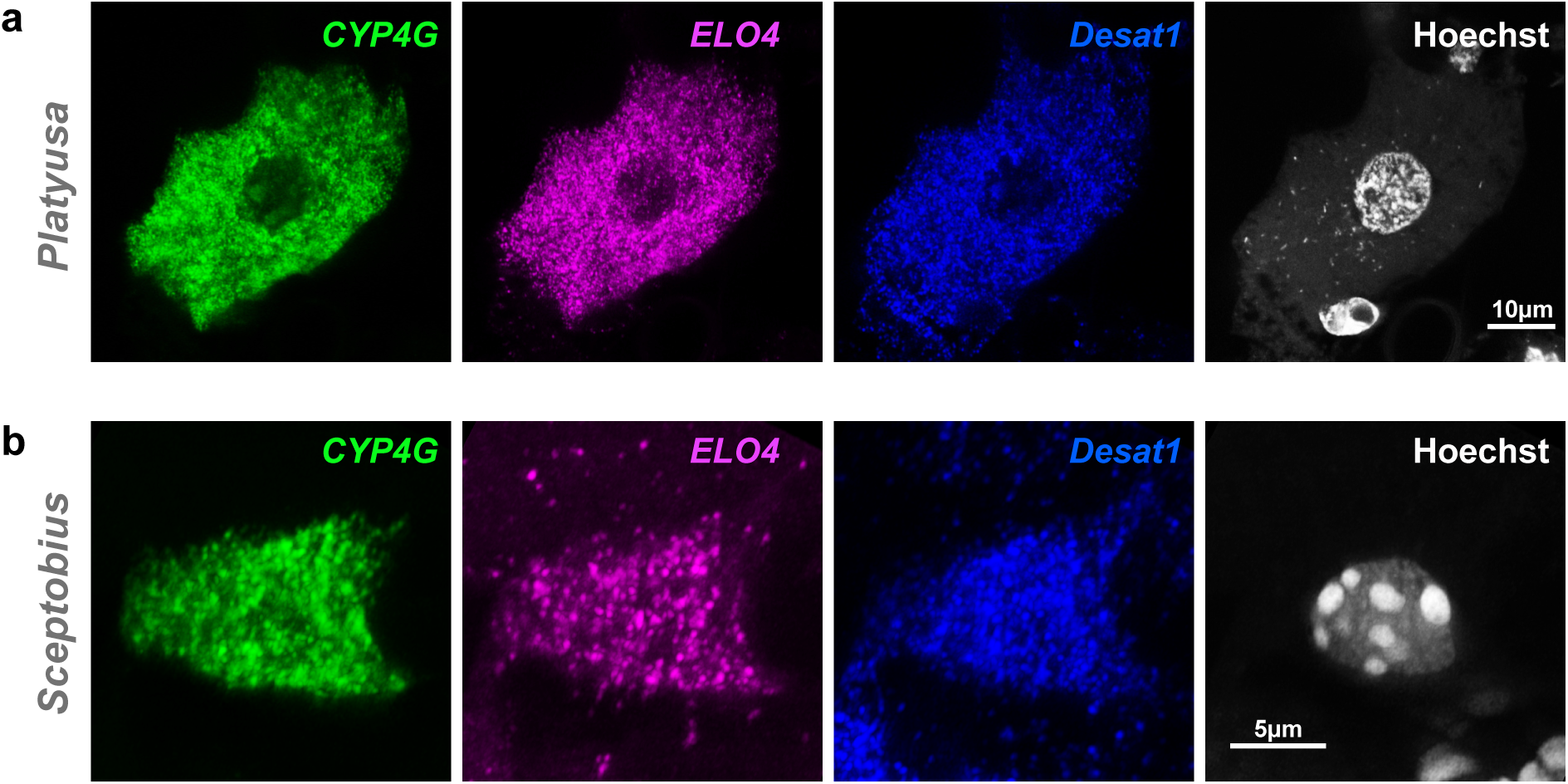
Additional CHC enzyme expression. HCR labelling of *CYP4G* (green), *ELO4* (magenta), *Desat1* (blue), and Hoechst (white) in **a:** *Platyusa sonomae* and **b:** *Sceptobius lativentris*.

**Figure S5.**
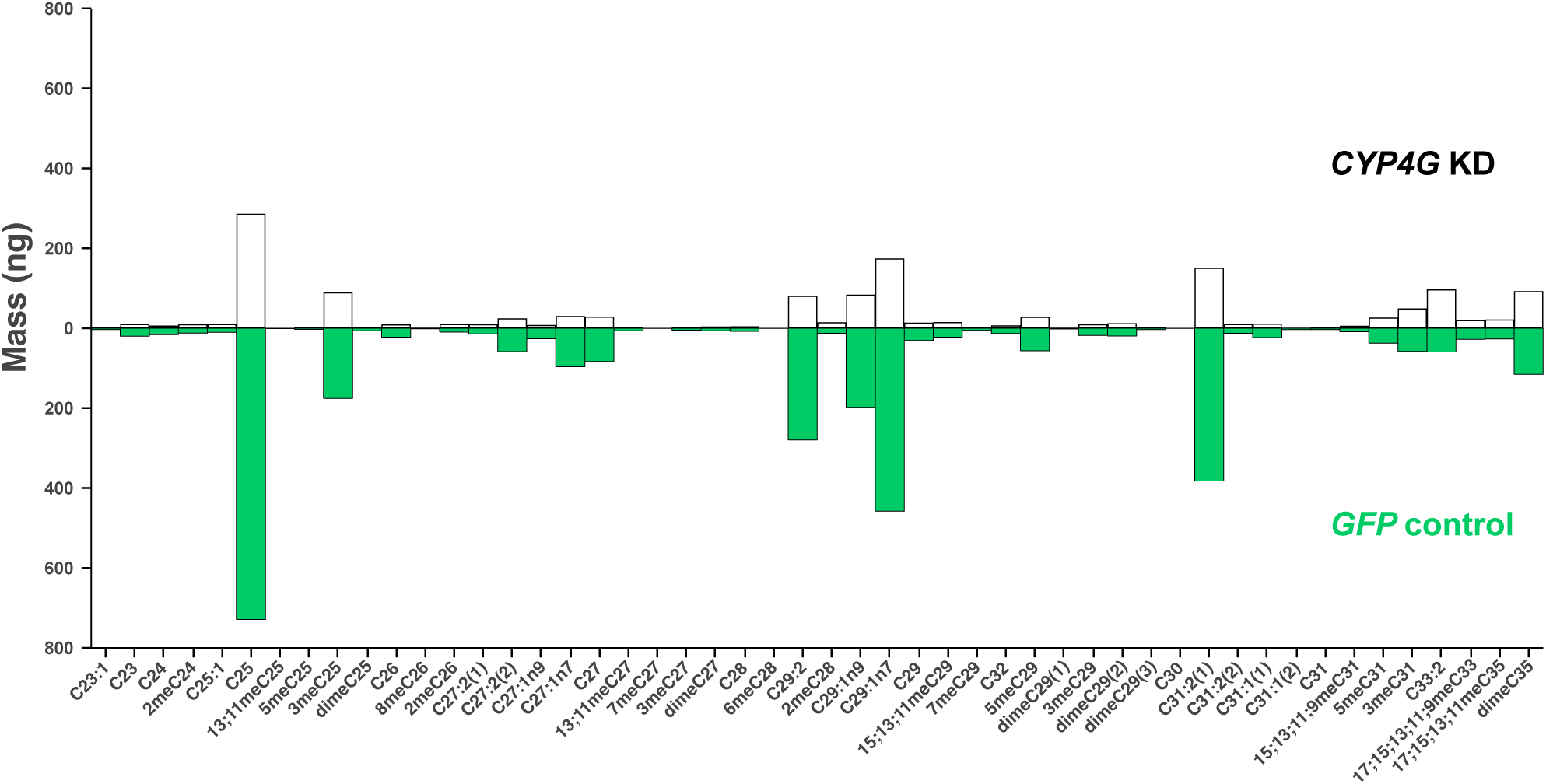
*CYP4G* RNAi in *Platyusa sonomae*. Average mass of each CHC extracted from wild caught *Platyusa sonomae* ten days after injecting with dsRNA targeting *CYP4G* or *GFP*.

**Figure S6.**
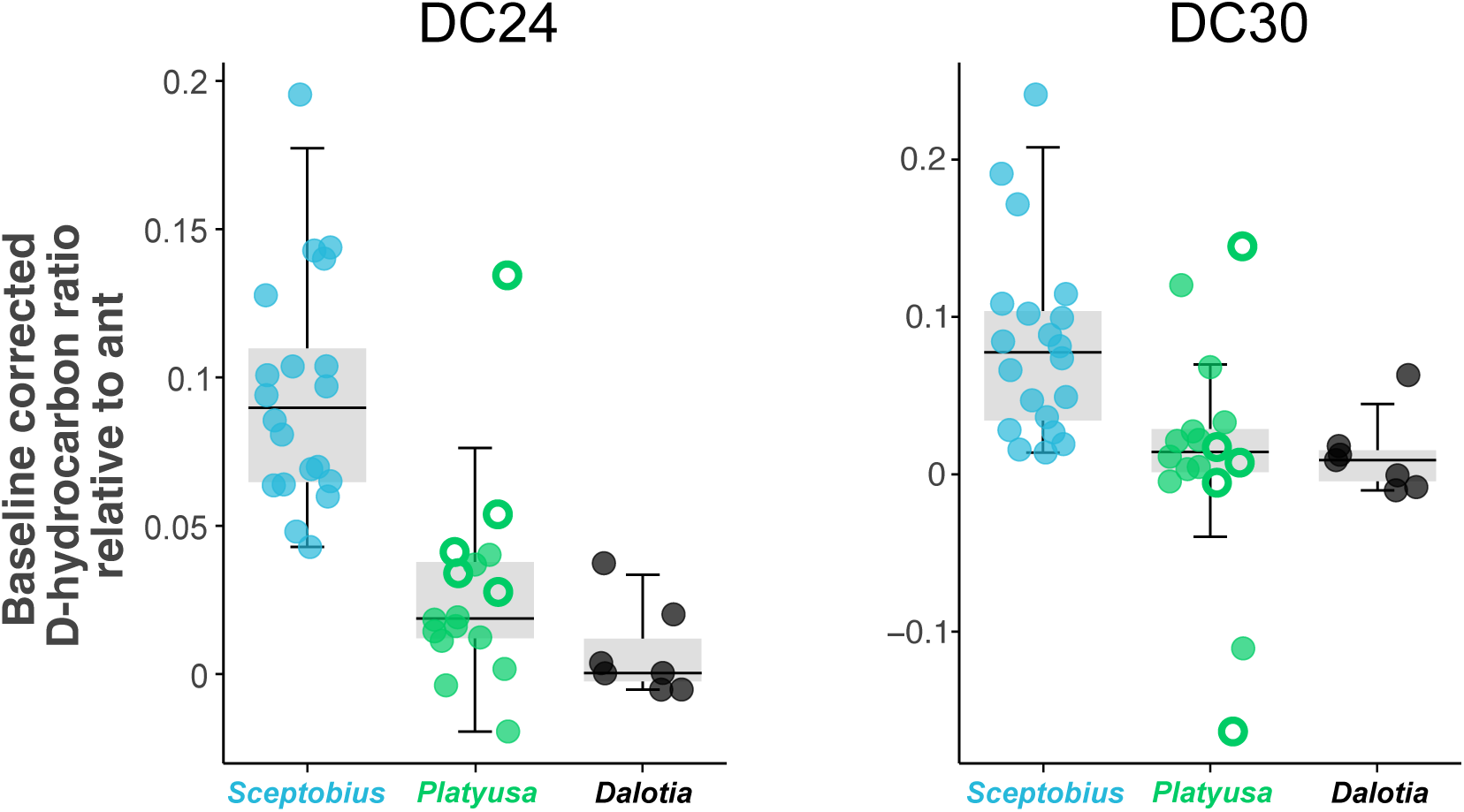
Additional Deuterated hydrocarbon transfer. Baseline-corrected deuterated hydrocarbon transferred from *Liometopum* to beetles during 24 hour interaction period using tetracosane-d_50_ (DC24) and triacontane-d_62_ (DC30). Open circles represent trials in which *Platyusa* ate the paired ant.

**Figure S7.**
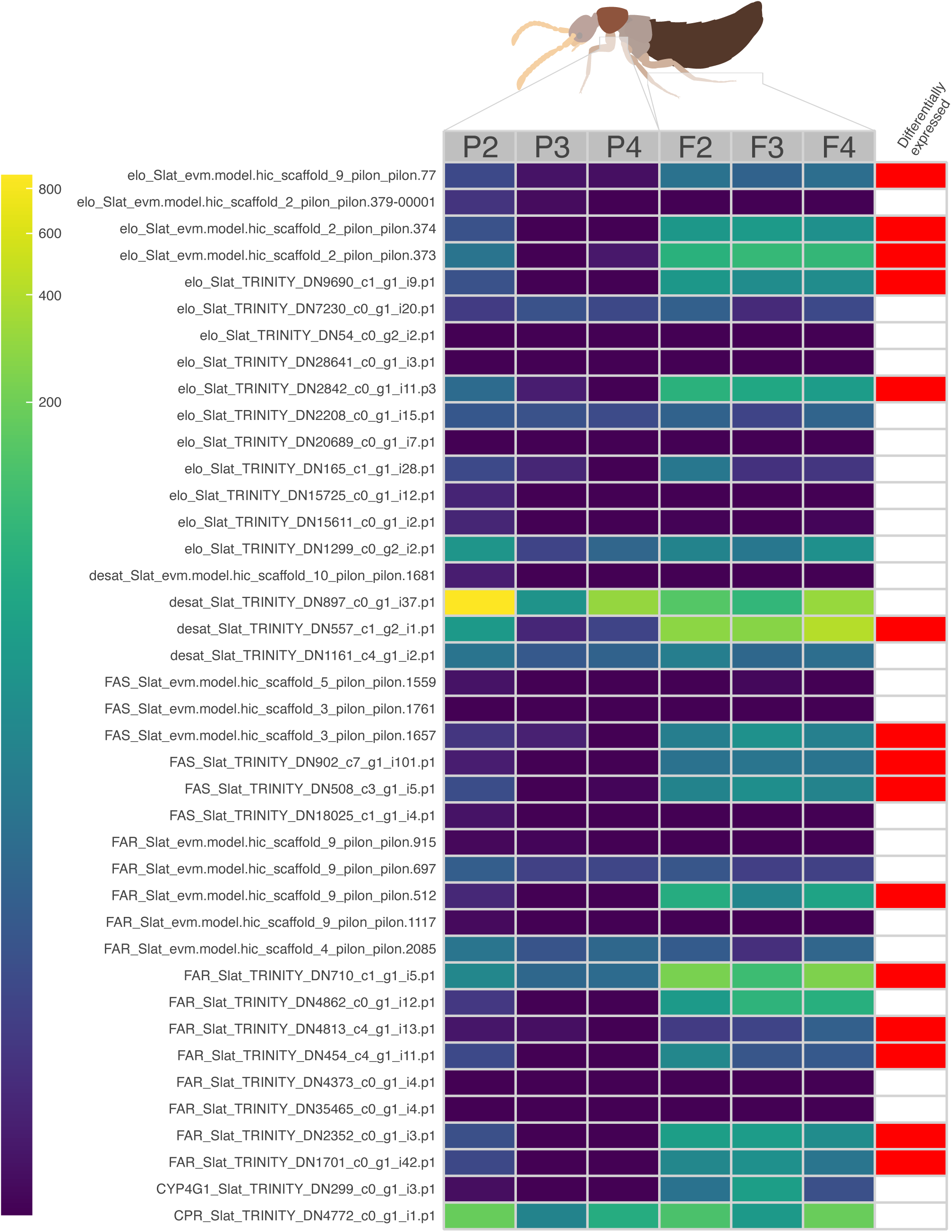
CHC biosynthesis enzyme expression in *Sceptobius lativentris*. Heatmap of CHC enzyme expression, in transcripts per million (TPM) for control tissue (pronotum, P) and test tissue (fat body, F). Numbers next to the treatment group correspond to the id of the single beetle dissected for that sample. Differentially expressed transcripts are shown in red.

**Figure S8.**
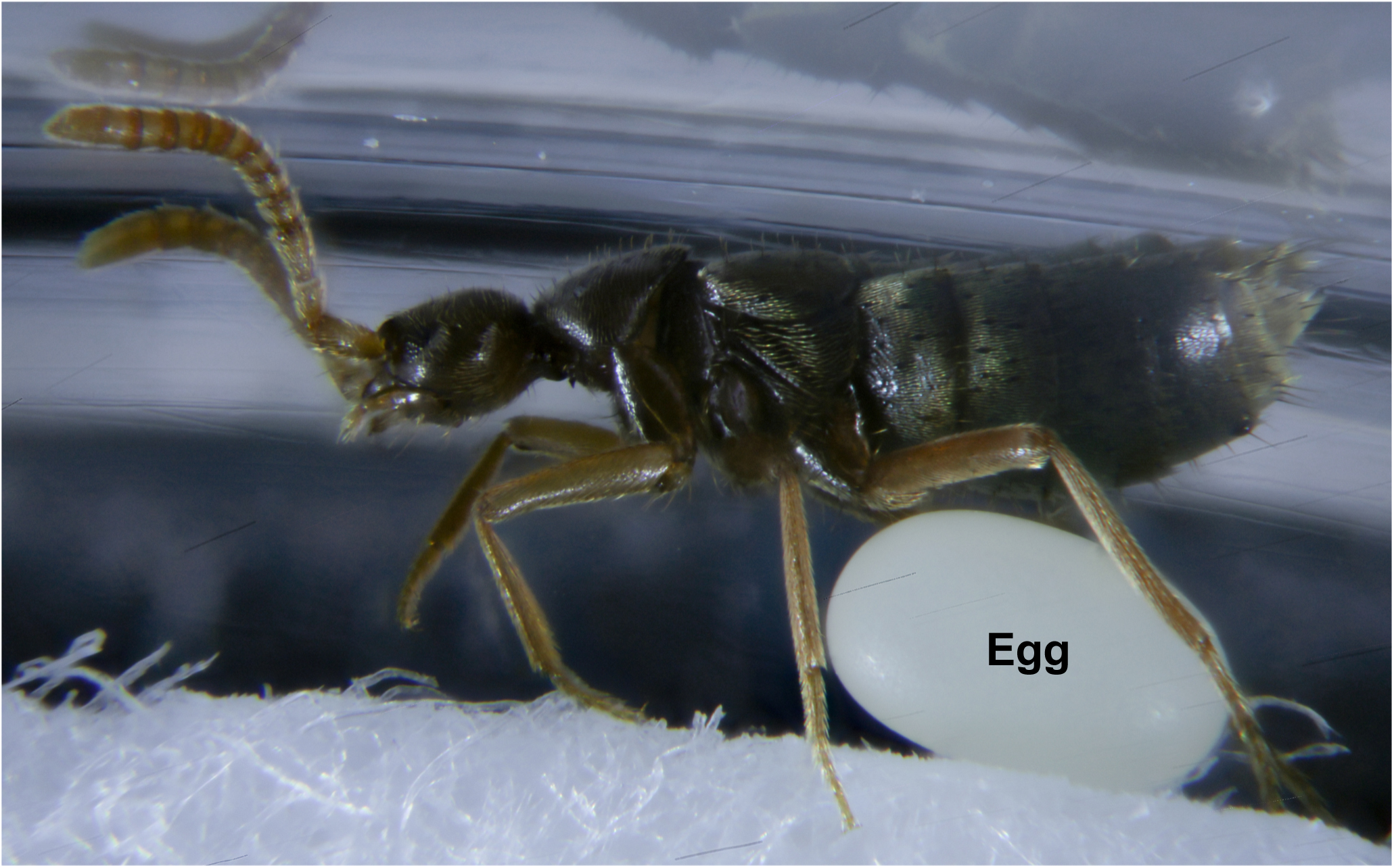
*Sceptobius* invest heavily in individual egg production. A deceased female *Sceptobius* next to her recently laid egg.

**Figure S9.**
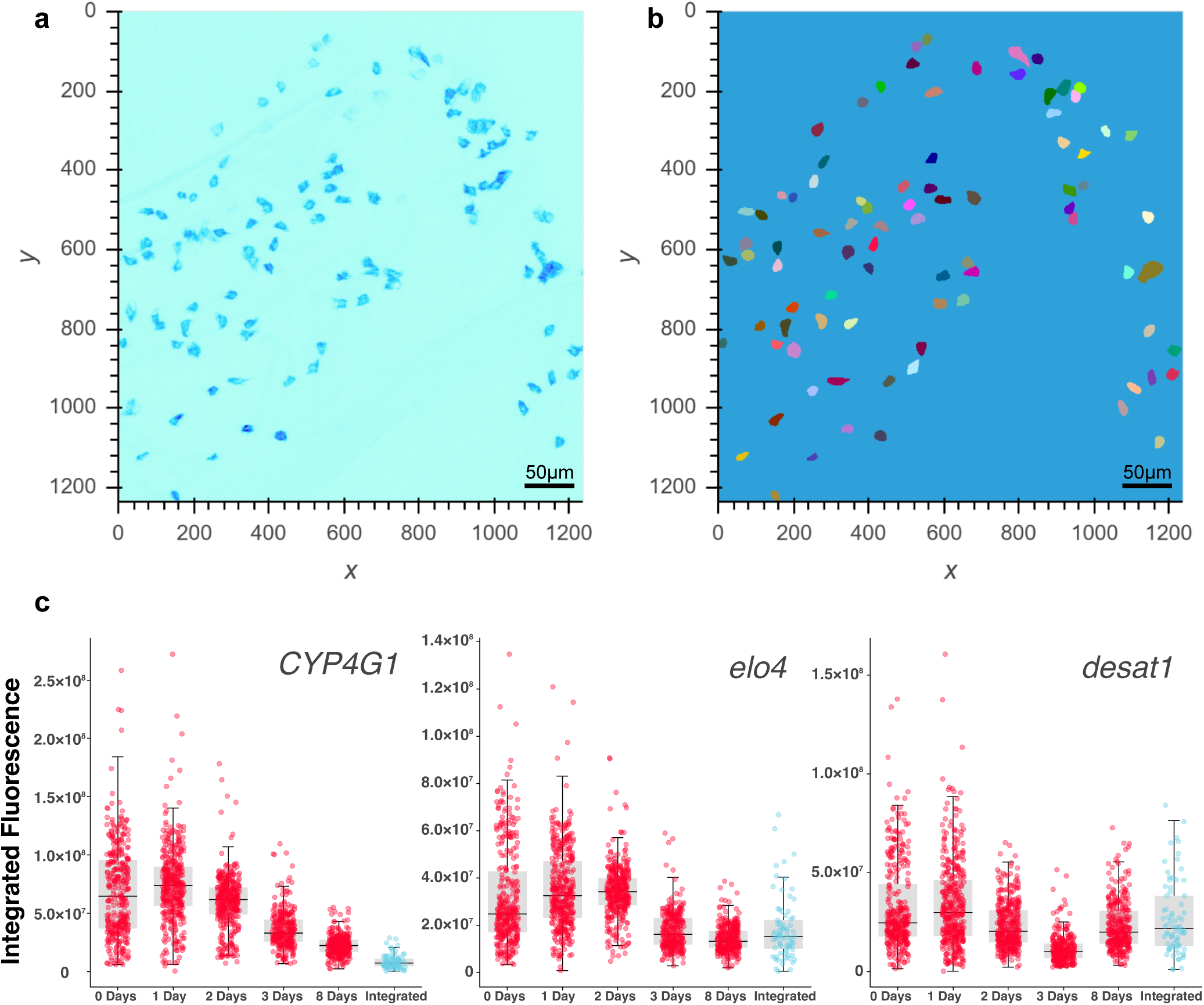
Cellpose analysis of CHC enzyme expression in *Sceptobius* post-eclosion. **a:** Summed-projection of a representative z-stack of HCR targeting *CYP4G* expression in teneral *Sceptobius*. **b:** Cellpose cell masks generated for the same sample. **c:** integrated fluorescence intensity of HCR probes targeting *CYP4G, ELO4*, and *Desat1* in individual oenocytes at multiple timepoints after *Sceptobius* eclosion.

**Figure S10.**
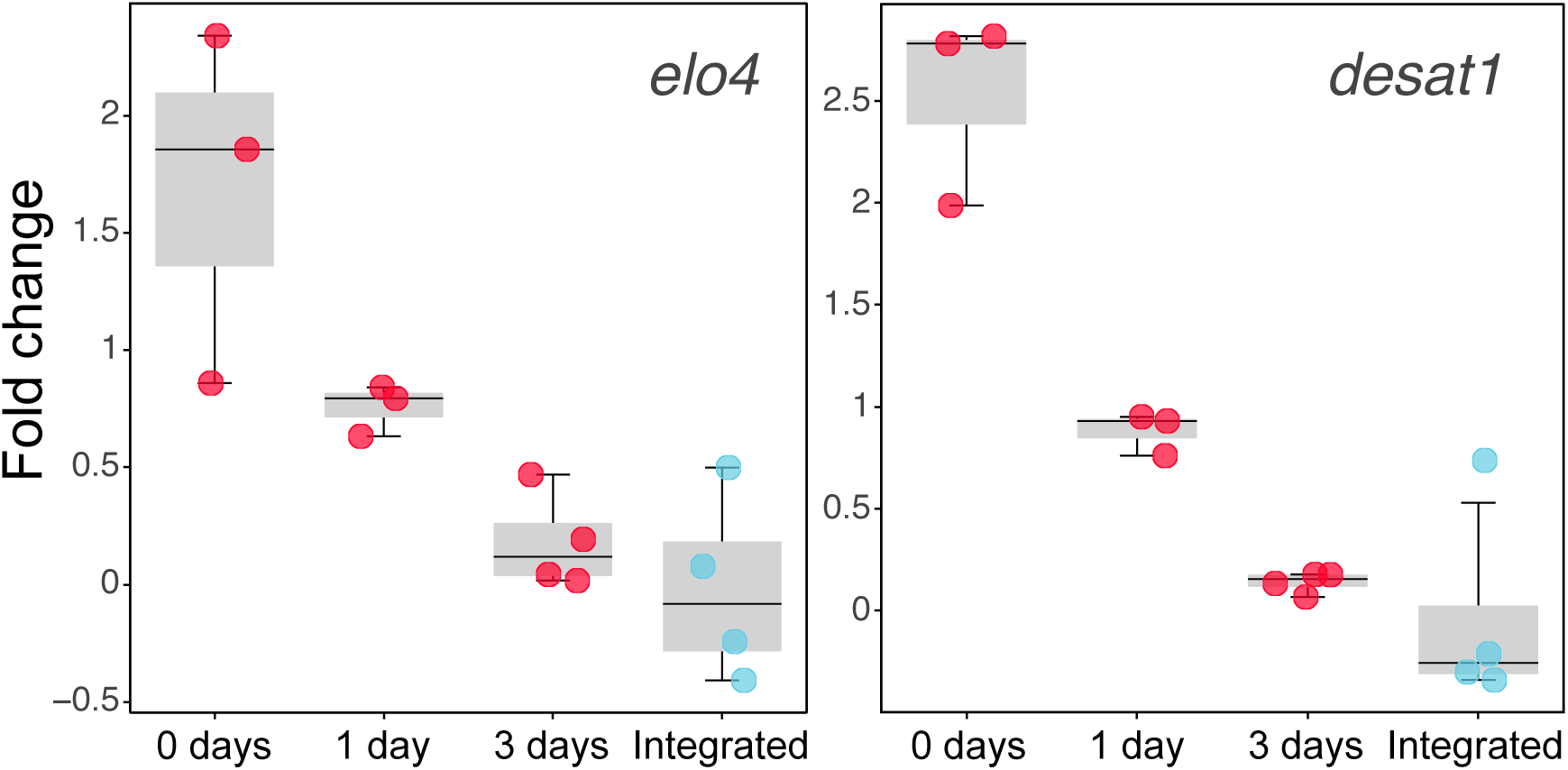
Additional qPCR measurements of CHC enzyme expression in *Sceptobius* post-eclosion. qPCR measurements of *ELO4* and *Desat1* transcription in whole-body extractions of *Sceptobius* at multiple time points following eclosion.

**Figure S11.**
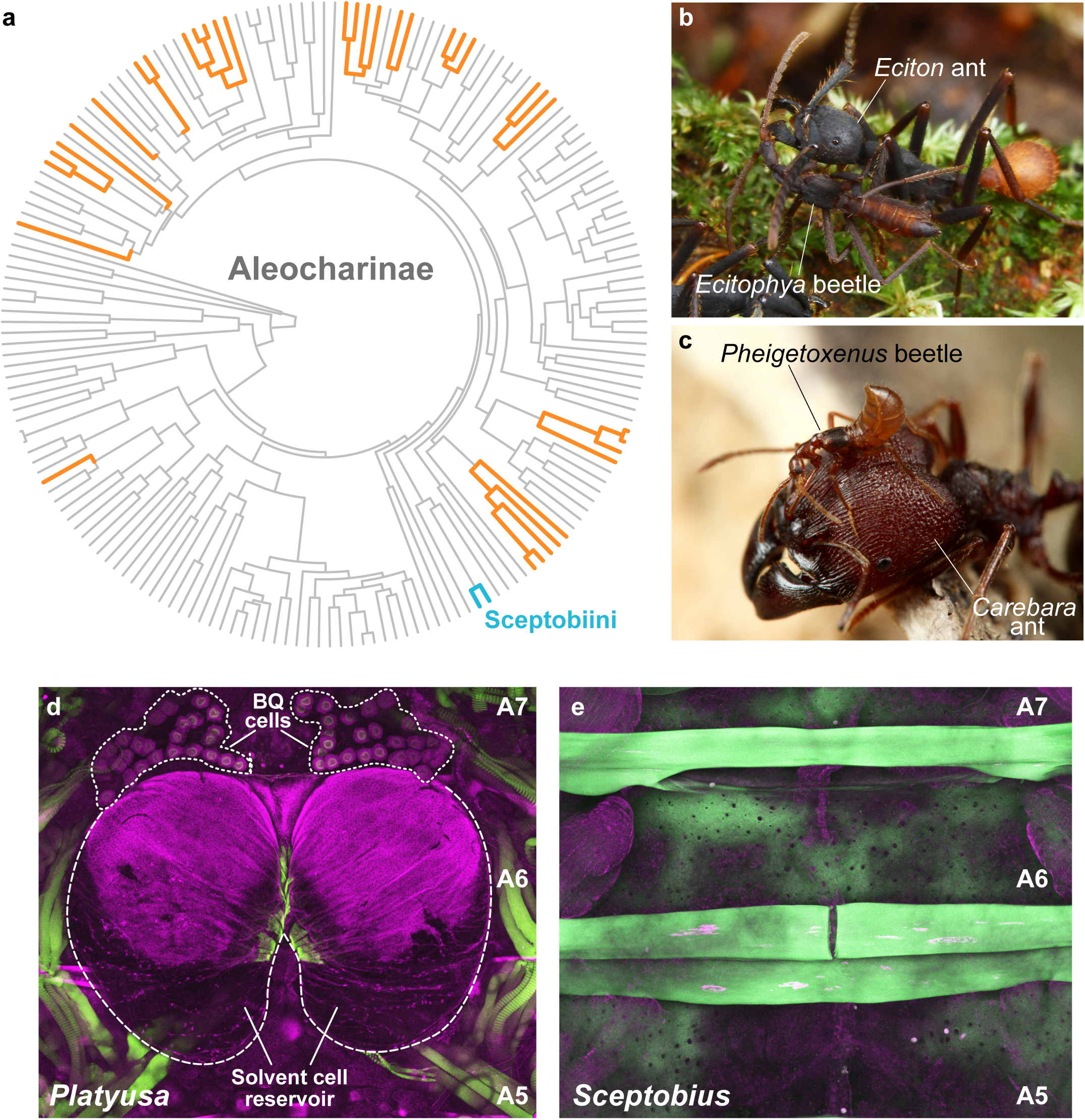
The Catch-22 syndrome across Aleocharinae. **a.** Aleocharinae phylogeny showing socially integrated clades in orange, or blue for Sceptobiini. Within these clades, no instances of reversion to a free-living lifestyle have been described. **b, c:** Examples of aleocharine myrmecophiles that have converged on a similar, obligately symbiotic phenotype to *Sceptobius*. **b.** *Ecitophya* grooming its host ant, *Eciton burchellii.* Credit: T. Komatsu **c.** *Pheigetoxenus* on a major worker of its host ant, *Carabera diversus*. Credit: Taku Shimada. **d:** The defensive tergal gland in *Platyusa sonomae,* which is present in all higher Aleocharinae (magenta, engrailed antibody and autofluorescence, general structures; green, phalloidin, muscles). The gland comprises two cell types: benzoquinone-secreting BQ cells, and alkane/ester secreting solvent cells, which form a large chemical reservoir inside the body cavity^64,65,161^. A5-A7 represent abdominal segments 5 to 7. **e.** The tergal gland has been evolutionarily lost in *Sceptobius lativentris* (green, autofluorescence; magenta, engrailed antibody and autofluorescence). Numbers represent abdominal segments. The gland is known to have been convergently lost in most or all of the orange clades in panel **a**^93–96,162–165^.

